# Transposable element profiles reveal cell line identity and loss of heterozygosity in *Drosophila* cell culture

**DOI:** 10.1101/2021.04.24.441253

**Authors:** Shunhua Han, Preston J. Basting, Guilherme Dias, Arthur Luhur, Andrew C. Zelhof, Casey M. Bergman

## Abstract

Cell culture systems allow key insights into biological mechanisms yet suffer from irreproducible outcomes in part because of cross-contamination or mislabelling of cell lines. Cell line misidentification can be mitigated by the use of genotyping protocols, which have been developed for human cell lines but are lacking for many important model species. Here we leverage the classical observation that transposable elements (TEs) proliferate in cultured *Drosophila* cells to demonstrate that genome-wide TE insertion profiles can reveal the identity and provenance of *Drosophila* cell lines. We identify multiple cases where TE profiles clarify the origin of *Drosophila* cell lines (Sg4, mbn2, and OSS_E) relative to published reports, and also provide evidence that insertions from only a subset of LTR retrotransposon families are necessary to mark *Drosophila* cell line identity. We also develop a new bioinformatics approach to detect TE insertions and estimate intra-sample allele frequencies in legacy whole-genome shotgun sequencing data (called ngs_te_mapper2), which revealed copy-neutral loss of heterozygosity as a mechanism shaping the unique TE profiles that identify *Drosophila* cell lines. Our work contributes to the general understanding of the forces impacting metazoan genomes as they evolve in cell culture and paves the way for high-throughput protocols that use TE insertions to authenticate cell lines in *Drosophila* and other organisms.

## Introduction

Cultured cell lines play essential roles in biological research, providing model systems to support discovery of basic molecular mechanisms and tools to produce biomolecules with medical and industrial relevance. Despite their widespread use, experiments in cultured cells often show non-reproducible outcomes, and increasing the rigor of cell-line based research is a priority of both funders and journals alike (Lorsch *et al.* 2014). One major source of irreproducible research comes from mislabelling or cross-contamination of cell lines (collectively referred to here as “misidentification”), resulting in cells of the wrong type or species being used in a particular study (Defendi *et al.* 1960; Gartler 1967; Nelson-Rees *et al.* 1981; MacLeod *et al.* 1999; Huang *et al.* 2017). As such, substantial effort has been invested into minimizing cell line misidentification through genotyping cell lines, cataloguing misidentified lines, standardizing cell line nomenclature, and the use of research resource identifiers (Masters *et al.* 2001; Capes-Davis *et al.* 2010; Barallon *et al.* 2010; Yu *et al.* 2015; Babic *et al.* 2019).

Starting with the first reports on the cell line misidentification problem, a variety of cytological and molecular techniques have been developed to authenticate mammalian cell lines (Defendi *et al.* 1960; Gartler 1967; O’Brien *et al.* 1977; Gilbert *et al.* 1990; Masters *et al.* 2001; Castro *et al*. 2013). These efforts culminated in development of short tandem repeats (STRs) as a widely-used standard to authenticate human cell lines at the molecular level (Masters *et al.* 2001; Barallon *et al.* 2010; Almeida *et al.* 2016). STR-based authentication has mitigated – but not eradicated – the human cell line misidentification problem, in part because of limitations in the stability, measurement, and matching of STRs (Parson *et al.* 2005; American Type Culture Collection Standards Development Organization Workgroup ASN-0002 2010; Yu *et al*. 2015; Horbach and Halffman 2017). More recently, alternative methods for genotyping human cell lines based on single nucleotide polymorphisms (SNPs) have been developed (Castro *et al.* 2013; Yu *et al.* 2015; Liang-Chu *et al.* 2015; Zaaijer *et al.* 2017; Mohammad *et al.* 2019), but these methods have not yet been accepted as standards for cell line authentication in humans (Almeida *et al.* 2016).

For most species beside humans, cell line authentication standards and protocols remain to be established (Almeida *et al.* 2016). For example, no protocols currently exist to authenticate cell lines in the fruitfly *Drosophila melanogaster*, despite the existence of over 150 different cell lines for this model animal system (Luhur *et al*. 2019). As such, no evidence of misidentified *Drosophila* cell lines have been catalogued to date by the International Cell Line Authentication Committee (v10, https://iclac.org/databases/cross-contaminations/). Development of cell line identification protocols and standards for common model organisms like *Drosophila* is an important goal for increasing rigor and reproducibility in bioscience. Achieving this goal for a new species requires an understanding of the genome biology and cell line diversity of that organism, and should ideally take advantage of powerful, cost-effective modern genomic technologies.

Relative to humans, the STR mutation rate is low in *D. melanogaster* (Schug *et al.* 1997) and thus the use of STRs for discriminating different *Drosophila* cell lines is likely to be limited. In contrast, it is well-established that transposable element (TE) insertions are highly polymorphic among individual flies (Charlesworth and Langley 1989), that TE abundance is elevated in *Drosophila* cell lines (Potter *et al.* 1979; Ilyin *et al.* 1980), and that TE families amplified in cell culture vary among *Drosophila* cell lines (Echalier 1997). These properties suggest that TE insertions should be useful markers to discriminate different cell lines established from distinct *D. melanogaster* donor genotypes (e.g. S2 vs Kc cells) and possibly also from the same donor genotype, including divergent sub-lines of the same cell line (e.g. S2 vs S2R+ cells) (Echalier and Ohanessian 1969; Schneider 1972; Yanagawa *et al.* 1998). Indeed, previous studies have shown that *D. melanogaster* cell lines have unique TE landscapes, and that sub-lines of the same cell line often share a higher proportion of TE insertions relative to distinct cell lines (Sytnikova *et al.* 2014; Rahman *et al.* 2015).

Here we show that *Drosophila* cell lines can successfully be clustered and identified on the basis of their genome-wide TE profiles using a combination of publicly available paired-end short-read whole genome shotgun (WGS) sequencing data from the modENCODE project (Lee *et al.* 2014) and new WGS data for eight widely-used *Drosophila* cell lines. Our approach reveals the first examples where the reported provenance of *Drosophila* cell lines – Sg4 (Morales *et al.* 2004) and mbn2 (Gateff *et al.* 1980) – conflicts with identity inferred from genomic data. Importantly, our TE-based clustering approach also allows us to identify which subset of TE families discriminate the most widely used *Drosophila* cell lines, paving the way for development of PCR-based genotyping protocols that can be used for cost-effective *Drosophila* cell line identification.

Additionally, we develop a new tool for detection of TEs in single-end whole genome shotgun data (called ‘ngs_te_mapper2‘) and integrate our new data with legacy data (Sienski *et al.* 2012; Sytnikova *et al.* 2014) to resolve the history and provenance of the widely-used OSS and OSC ovarian cell lines (Niki *et al.* 2006; Saito *et al.* 2009). Using TE-based clustering, we provide evidence that OSS and OSC cell lines can be discriminated on the basis of the ZAM retrotransposon family. We propose that the OSS_E sub-line reported in Sytnikova *et al*. (2014) approximates an ancestral state of the OSC cell line, with contemporary OSC sub-lines having undergone loss of heterozgosity (LOH) in cell culture from an OSS_E-like state. Together, our results show that TE insertions are a powerful source of genetic markers that can be used for cell line authentication in *Drosophila* and that LOH is an important mechanism driving *Drosophila* cell line genome evolution.

## Results and Discussion

### Clustering of cell lines using TE insertions reveals rare cases of mismatch with expected provenance

We reasoned that TE insertions would be favorable genetic markers for cell line identification in *Drosophila* because the joint processes of germline transposition in whole flies and somatic transposition in cell culture together would create unique TE profiles, both for cell lines derived from distinct *D. melanogaster* donor genotypes and for sub-lines of cells derived from the same original donor genotype (Fig 1). Furthermore, we posited that shared presence or absence of TE insertions at orthologous loci would allow the identity or similarity among cell line samples to be assessed based on a clustering approach.

**Figure 1.**
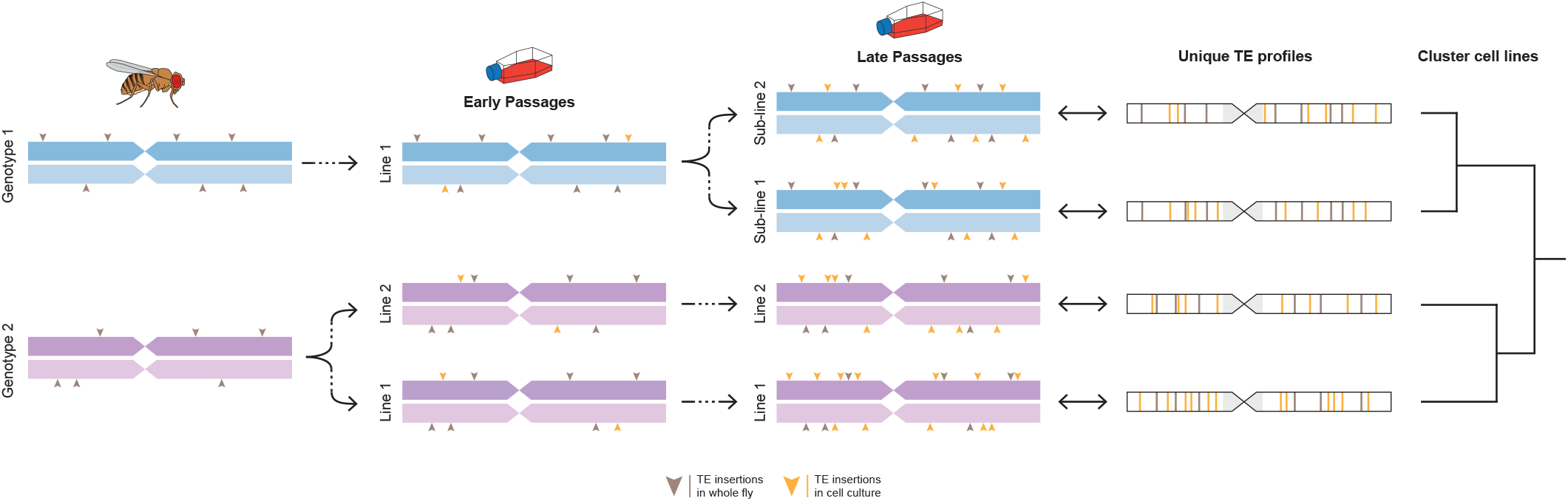
Germline and somatic transposition jointly can create unique TE profiles in *Drosophila* cell line genomes. A homologous pair of chromosomes is shown for two donor fly genotypes used to establish two distinct cell lines. TE profiles initially differ because transposition events in whole flies (grey arrowheads) are maintained at low population frequencies by purifying selection. After establishment of distinct cell lines, ongoing transposition in cell culture (orange arrowheads) further differentiates TE profiles, both for distinct cell lines derived from the same or different donor genotypes as well as for sub-lines of the same cell line. Ultimately these processes lead to unique TE profiles that can identify cell lines and allow them to be clustered based on shared presence or absence of TE insertions at orthologous loci. The model depicts a simplified case of diploidy, when in reality cell culture genomes can have complex genome structure due to polyploidy and segmental aneuploidy.

We initially investigated the possibility of TE-based cell line identification in *Drosophila* using public genome sequences for 26 samples from 18 cell lines generated by the modENCODE project (Lee *et al.* 2014) (Table S1). Paired-end Illumina WGS sequences were used to predict non-reference TEs using TEMP (Zhuang *et al.* 2014), which showed the least dependence on read length (Fig. S1) or coverage (Fig. S2) out of eight non-reference TE detection methods tested on the data used in this study. We clustered cell lines on the basis of their TE profiles using Dollo parsimony, which accounts for the virtually homoplasy-free nature of TE insertions within species (Batzer and Deininger 2002; Ray *et al.* 2006), the ancestral state of TE absence at individual loci (Batzer and Deininger 2002) and false negative predictions inherent in non-reference TE detection software (Nelson *et al*. 2017; Rishishwar *et al.* 2017; Vendrell-Mir *et al.* 2019). Use of Dollo parsimony for clustering cell line samples also allows ancestral states to be reconstructed, facilitating inference of which TE families diagnostically identify individual cell lines or groups of cell lines. We note that we do not attempt to interpret the clustering relationships among distinct cell lines in an evolutionary context, however our approach does provide insight into the evolutionary history of clonally-evolving sub-lines established from the same original cell line.

We predicted between 730 and 2579 non-reference TE insertions in euchromatic regions of *Drosophila* cell line samples from the modENCODE project (Table S2). As reported previously for human cancer cell lines (Zampella *et al.* 2016), each *Drosophila* cell line sample had a unique profile of TE insertions (File S1). The most parsimonious clustering of *Drosophila* cell lines using TE profiles revealed several expected patterns that indicate TE insertions reliably mark the identity of *Drosophila* cell lines (Fig. 2A, File S2). First, replicate samples of the same cell line cluster most closely with one another with 100% bootstrap support in all seven cases where data is available (S2, S2R+, CME-W1-Cl.8+, ML-DmD9, ML-DmD16-c3, ML-DmD20-c5, and Kc167). Second, different cell lines created in the same lab (presumably from the same ancestral fly genotype) cluster with each other before they cluster with cell lines generated in other labs, or with cells lines having different ancestral genotypes. Third, we observe that divergent sub-lineages of the same cell line (i.e. S2 and S2R+) cluster closely together (Schneider 1972; Yanagawa *et al*. 1998). We also find weak evidence for clustering of cell lines generated in different labs (Schneider, Milner) that are derived from the same putative ancestral fly stock (Oregon-R). However, we caution against over-interpretation of this result, given previous reports for substantial genetic diversity among common lab stocks like Oregon-R (Rahman *et al.* 2015; Stanley and Kulathinal 2016). Also, cell lines derived from the Schneider and Milner labs have distinct B-allele frequency (BAF) profiles, suggesting different ancestral Oregon-R genotypes (Fig. S3B).

**Figure 2.**
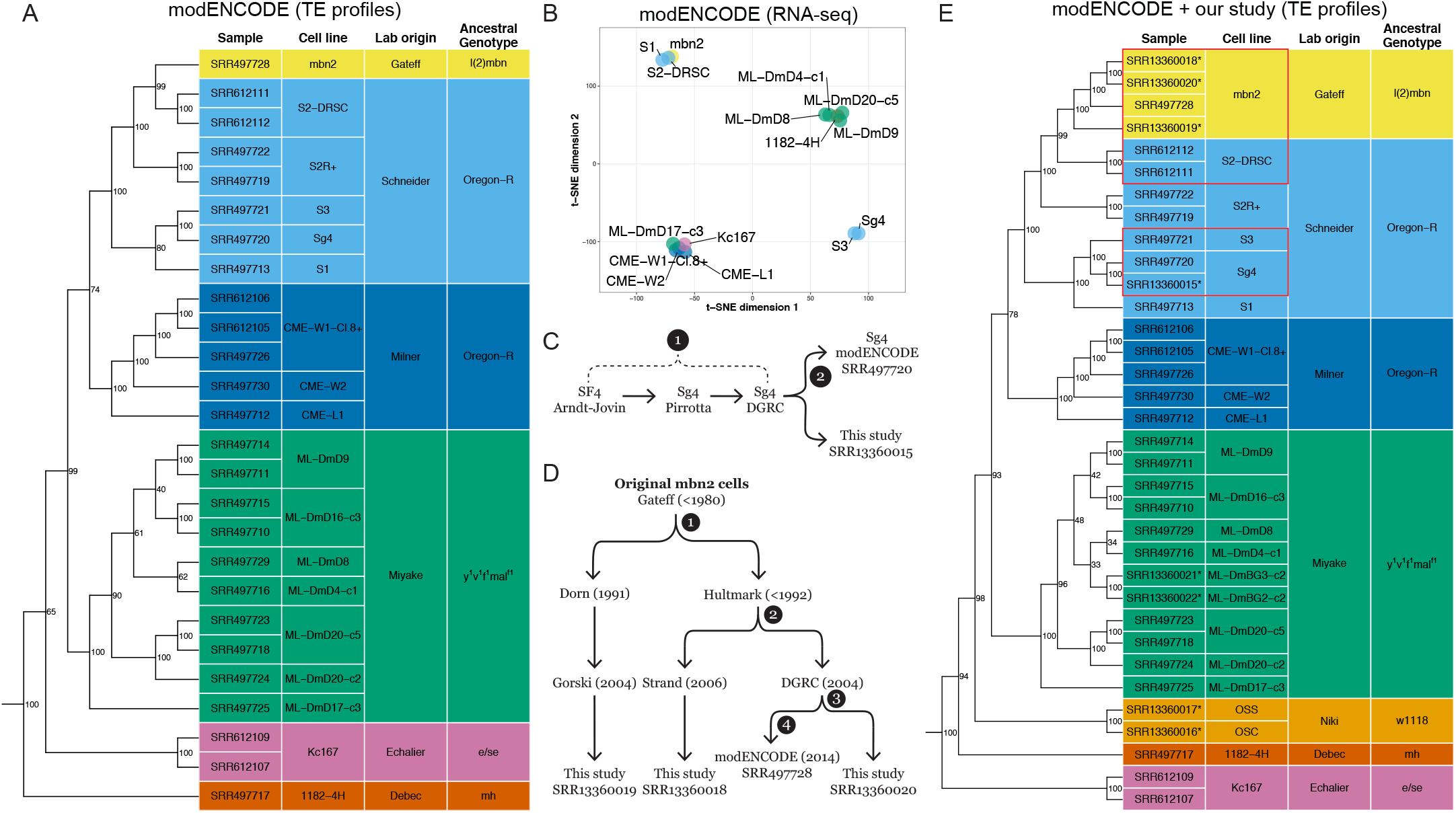
TE insertion profiles cluster *Drosophila* cell lines by lab origin and reveal unexpected placement of the Sg4 and mbn2 cell lines. (A) Clustering of *Drosophila* cell line samples from the modENCODE project was constructed using Dollo parsimony based on non-reference TE insertions. Samples are colorized by the lab origin based on the first publication reporting the original variant of the cell line. Ancestral genotype is based on the *D. melanogaster* stock reported to create the original variant of the cell line. (B) t-SNE visualization of 15 *Drosophila* cell line samples using transcriptomic data in (Stoiber *et al.* 2016). Samples are colorized by the lab origin of cell lines. (C) Key events in the history of the Sg4 cell line creation and distribution. (D) Key events in the history of the mbn2 cell line distribution. Node labels in panels C and D represent timepoints in the past that potential cell line misidentification events could have occurred. (E) Clustering of *Drosophila* cell line samples from the modENCODE project plus new data reported here (indicated by asterisks in panel E) was constructed using Dollo parsimony based on non-reference TE insertions. Numbers beside nodes in panels A and E indicate percent support based on 100 bootstrap replicates. Red boxes in panel E highlight cases where the reported provenance of *Drosophila* cell lines conflicts with identity inferred from genomic data.

Overall, clustering patterns based on TE profiles suggest that misidentification is rare among the panel of cell lines sequenced by modENCODE. However, we observed two cases where the similarity of cell lines based on genome-wide TE profiles conflicted with expectations based on reported provenance. First, we unexpectedly found that the Sg4 cell line (originally called Sf4 by its maker Donna Arndt-Jovin) clusters most closely with S3 cells, although the DGRC and FlyBase currently consider Sg4 to be a variant of S2 cells (http://flybase.org/reports/FBrf0205934.html; http://flybase.org/reports/FBtc0000179; https://dgrc.bio.indiana.edu/cells/S2Isolates). More strikingly, we also observed that the mbn2 cell line originally reported by Gateff *et al*. (1980) to be derived from the l(2)mbn stock was placed inside a well-supported cluster containing cell lines (S1, S2, S2R+, S3, Sg4) generated by Schneider (1972) from an Oregon-R stock. Our clustering of mbn2 cells inside the Schneider cell clade is consistent with a previously unexplained observation that mbn2 cells share an unexpectedly high proportion of TE insertions with both S2 and S2R+ cells (Rahman *et al.* 2015).

Clarification of the provenance of the Sg4 and mbn2 cell lines used by modENCODE is important since many functional genomics resources were generated for these cell lines (Roy *et al.* 2010) and over 125 publications involving these cell lines are curated in FlyBase (Larkin *et al.* 2021). To cross-validate genomic clustering based on TE profiles and to assess potential functional similarity between Sg4↔S3 and mbn2↔S2 cell lines, we clustered cell lines on the basis of their transcriptomes. Transcriptome-based clustering should reveal similarities among cell types rather than genotypes, and thus is not expected to globally match our TE insertion based clustering. However, both cell type and genotype clustering should support the similarity of pairs of cell lines that are derived from a common ancestral cell line.

Previous transcriptome-based clustering of cell lines based on early whole-genome tiling microarray datasets from the modENCODE project did not reveal similarities among Sg4 and S3 or mbn2 and S2 (Cherbas *et al.* 2011), however clustering of small RNA-seq data did reveal similarities among these cell lines (Wen *et al.* 2014). Using a consistent batch of poly-A RNA-seq samples from a panel of 15 DGRC cells lines with genome data (Stoiber *et al.* 2016) (Table S3), we estimated expression levels for protein-coding genes then used T-distributed Stochastic Neighbor Embedding (t-SNE) dimensionality reduction (Maaten and Hinton 2008; Maaten 2014) to visualize similarity of cell lines based on their gene expression profiles. This analysis revealed that gene expression profiles based on transcriptome data support the clustering of Sg4 with S3 and mbn2 with S2 (Fig. 2B). Transcriptome-based clustering of Sg4 with S3 and mbn2 with S2 is also observed in a different batch of RNA-seq samples generated independently by the modENCODE project (Brown *et al*. 2014) (Fig. S4, Table S3). These results provide replicated transcriptomic support for the clustering of Sg4↔S3 and mbn2↔S2 cell lines revealed by TE profiles, and also highlight functional similarities between these pairs of cell lines.

### TE profiles help resolve the provenance of the Sg4 and mbn2 cell lines

To better understand the cause of the surprising patterns of clustering for the Sg4 and mbn2 cell lines in the modeENCODE data, we generated paired-end Illumina WGS sequences for additional samples of Sg4 and mbn2 cells from the DGRC and other sources. In addition, we sequenced several other popular *Drosophila* cell lines (OSS, OSC, ML-DmBG3-c2, ML-DmBG2-c2) that were not originally sequenced in the modENCODE cell line genome project (Lee *et al.* 2014). To guide sampling and aid the interpretation of the expanded dataset, we reconstructed key events in the history of the Sg4 (Fig. 2C) and mbn2 cell lines (Fig. 2D). We predicted non-reference TE insertions in these additional samples and then reclustered the expanded dataset using the same methods as the modENCODE-only dataset. Inclusion of additional samples altered some details of the clustering relationships among D-series cell lines generated by the Miyake lab and the position of distantly related cell lines with respect to the root (Kc167 and 1182-4H) (Fig. 2A vs E). However, key aspects of our clustering approach that facilitate cell line identification (replicates clustering most closely, clustering of cell lines from the same lab/ancestral genotype) appear to be robust to the set of cell line samples analyzed.

Clustering TE profiles from this expanded dataset of 34 samples from 22 *Drosophila* cell lines revealed that our resequenced sample of DGRC Sg4 clusters with high support first with the modENCODE sample of DGRC Sg4 then with S3 (Fig. 2E). This result confirms the reproducibility of the S3↔Sg4 genomic similarity and rejects the possibility of cell line swap during the modENCODE cell line sequencing project (node 2; Fig. 2C). Additional evidence for the similarity of Sg4 and S3 can be observed in their BAF and CNV profiles. All Sg4 and S3 samples are generally devoid of heterozygosity across their entire genomes, including lacking a small patch of heterozygosity at the base of chromosome arm 2L that is present in all S2 or S2R+ samples (Fig. S3B). All Sg4 and S3 samples also share CNVs on chromosome arms 2L and 3L that are not present in any S2/S2R+ sample (Fig. S3C). Together, these data support the conclusion that DGRC Sg4 is a variant of the S3 cell line, not the S2 cell line as currently thought. Presently, we are unable to determine where misidentification of Sg4 as a variant of S2 occurred in the provenance chain from initial development of the cell line by the Arndt-Jovin lab to receipt by the DGRC (node 1; Fig. 2C). Future analysis of additional Sg4 sub-lines circulating in the research community (Morales *et al.* 2004; Schwartz *et al.* 2006) will be necessary to establish the timing of this event and if the S3↔Sg4 similarity first observed in the DGRC Sg4 sub-line is more widespread.

The second case of unexpected clustering we observed in the modENCODE data involving mbn2 and S2 is more surprising and consequential given that these cell lines are reported to be derived from different ancestral genotypes. mbn2 cells were reportedly derived from a stock carrying l(2)mbn on a 2nd chromosome marked with three visible mutations (Gateff 1977; Gateff *et al.* 1980), while S2 cells were derived from a wild-type Oregon-R stock (Schneider 1972). Unfortunately, the l(2)mbn mutation was never characterized at the molecular level, and no fly stocks carrying l(2)mbn currently exist in public stock centers that could be sequenced and compared with the mbn2 cell line. In the absence of external biological resources to verify the identity of an authentic mbn2 cell line, we attempted to infer the timing and extent of the potential mbn2 misidentification event first observed in the modENCODE data by sequencing sub-lines of mbn2 from DGRC and other sources. We resequenced another sample of the DGRC mbn2 sub-line, a sub-line from the Strand lab (University of Georgia) derived from the same donor as the DGRC sub-line (Hultmark lab, Umeå University), and a sub-line from the Gorski lab (Canada’s Michael Smith Genome Sciences Centre, BC Cancer) derived from an independent donor (Dorn lab, Johannes Gutenberg-Universität Mainz) (Fig. 2D). The Hultmark and Dorn labs each report obtaining mbn2 cells directly from the Gateff lab in the early 1990s (Samakovlis *et al.* 1992; Ress *et al.* 2000). This sampling allowed us to infer if potential misidentification occurred during the modENCODE project (node 4), at the DGRC (node 3), in the Hultmark lab (node 2) or in the Gateff lab (node 1) (Fig. 2D).

Analysis of TE profiles in our expanded dataset revealed that all four samples of mbn2 cluster together as a single, well-supported group that is most similar to a cluster containing S2 cells (Fig. 2E). The detailed relationships among sub-lines within the mbn2 cluster deviate slightly from expectations based on cell line history (Fig. 2D), however this discrepancy appears to be caused by differences in read length or coverage between the data from modENCODE and our study (Fig. S5). All mbn2 samples have the low SNP heterozygosity across most of their genomes that is characteristic of Schneider cell lines, and also share the small patch of heterozygosity at the base of chromosome arm 2L found in S2 and S2R+ cells (Fig. S3B). Additionally, all four mbn2 samples share widespread segmental aneuploidy across the entire euchromatin that is a common hallmark of S2 and S2R+ cells, but not other *Drosophila* cell lines (Fig. S3C). Together, these data support the conclusions that multiple independent sub-lines of mbn2 cells all share a common origin and are likely to originally descend from a single divergent lineage of S2 cells. Based on these observations, we speculate that currently-circulating mbn2 cells derive from a mislabelling or cross-contamination event with S2 cells in the Gateff lab that occurred prior to distribution to the Hultmark or Dorn labs (node 4, Fig. 2D). This scenario is consistent with the facts that S2 cells were developed and widely distributed prior to the origin of mbn2 cells (Schneider 1972; Gateff *et al.* 1980) and that there was a 12 year gap between the initial report describing mbn2 cells and use in any subsequent publication (Gateff *et al.* 1980; Samakovlis *et al.* 1992).

The possibility that mbn2 cells are essentially a divergent lineage of S2 cells is plausible given that both cell lines are thought to have a hemocyte-like cell type (Cherbas *et al.* 2011; Luhur *et al.* 2019). Furthermore, it is known that different lineages of *bona fide* S2 cells vary substantially in their morphology and gene expression, some of which share properties with mbn2 cells (Samakovlis *et al.* 1992; Yanagawa *et al.* 1998; Cherbas *et al*. 2011) (Fig. S6). Under phase-contrast microscopy, canonical S2 cells represented by the S2-DRSC sub-line are generally a mix of loosely adherent spherical cells and simple round flat cells. In contrast, live S2R+ cells can be characterized by many “phase dark” cells that attach to the growth substrate, which can flatten out to exhibit both polygonal and “fried egg” morphology. S2R+ cells that are loosely attached to the growth surface are generally spherical with fine cell protrusions. Like S2R+ cells, mbn2 cells are characterized by a mix of flattened phase dark cells that assume the polygonal and fried egg morphology, as well as loosely adhering spherical cells. However, loosely adherent mbn2 cells have a bigger diameter relative to S2-DRSC and S2R+ cells. Recognition of mbn2 as a divergent S2 lineage suggests that complex morphology may be the ancestral state of all S2 lineages, and that there is more phenotypic diversity among different S2 lineages than previously recognized.

### A subset of LTR retrotransposon families are sufficient to identify Drosophila cell lines

Our analysis has thus far provided evidence that TE insertion profiles of commonly used *Drosophila* cell lines based on whole-genome sequences can be used to cluster cell lines and uncover cases of cell line misidentification. However, for these results to form the foundation for a *Drosophila* cell line authentication protocol, it is necessary to show that a cell line sample can successfully be identified on the basis of its TE profile. Furthermore, it is important to explore if whole-genome data is required for TE-based cell line identification in *Drosophila* since the cost of WGS could preclude its routine application by many labs. Therefore, we next investigated whether a subset of *Drosophila* TE families could potentially be sufficient for *Drosophila* cell line identification, with the aim of guiding development of a cost-effective targeted PCR-based enrichment protocol that could be used more widely by the research community.

To investigate this possibility, we first clustered a non-redundant dataset of one “primary” replicate from each of the 22 *Drosophila* cell lines in the expanded dataset based on their whole-genome TE profiles (Fig. 3A), which resulted in a similar clustering to the same sample of 22 cell lines including all replicates (Fig. 2E). Replicates with the longest read length or depth of coverage were chosen as the primary replicate in the non-redundant dataset (Table S1). We then took advantage of the ability of Dollo parsimony to reconstruct ancestral states and map the gain of TE insertions on each branch of the most parsimonious tree. TE insertions were then aggregated into families on each branch of the tree to visualize family- and branch-specific TE insertion profiles. This analysis revealed that a subset of 60 out of the 125 curated TE families in *D. melanogaster* are informative for *Drosophila* cell line clustering using TEMP predictions (Fig. 3B, File S3). Within the set of clustering-informative TE families, we observed that some TE families are broadly represented across many cell lines with different origins (e.g. copia, 297, jockey, mdg3, mdg1, and roo), although the quantitative abundance of these TE families varies across cell lines. Other TE families appear to be represented in only one cell line or a subset of cell lines from the same lab origin (e.g. ZAM, Tabor, HMS-Beagle2, gypsy5, 1731, 17.6, springer, Tirant, rover, micropia). These results provide systematic genome-wide evidence for the classical observation that proliferation of different TE families in cultured *Drosophila* cells is cell-line dependent (Echalier 1997). Additionally, these patterns of cell-line specific TE proliferation provide further support for the conclusions that the DGRC Sg4 cell line is a lineage of S3 cells (all share Ivk proliferation), and that mbn2 cell lines are a divergent lineage of S2 cells (all share 1731 proliferation) (Fig. 3B).

**Figure 3.**
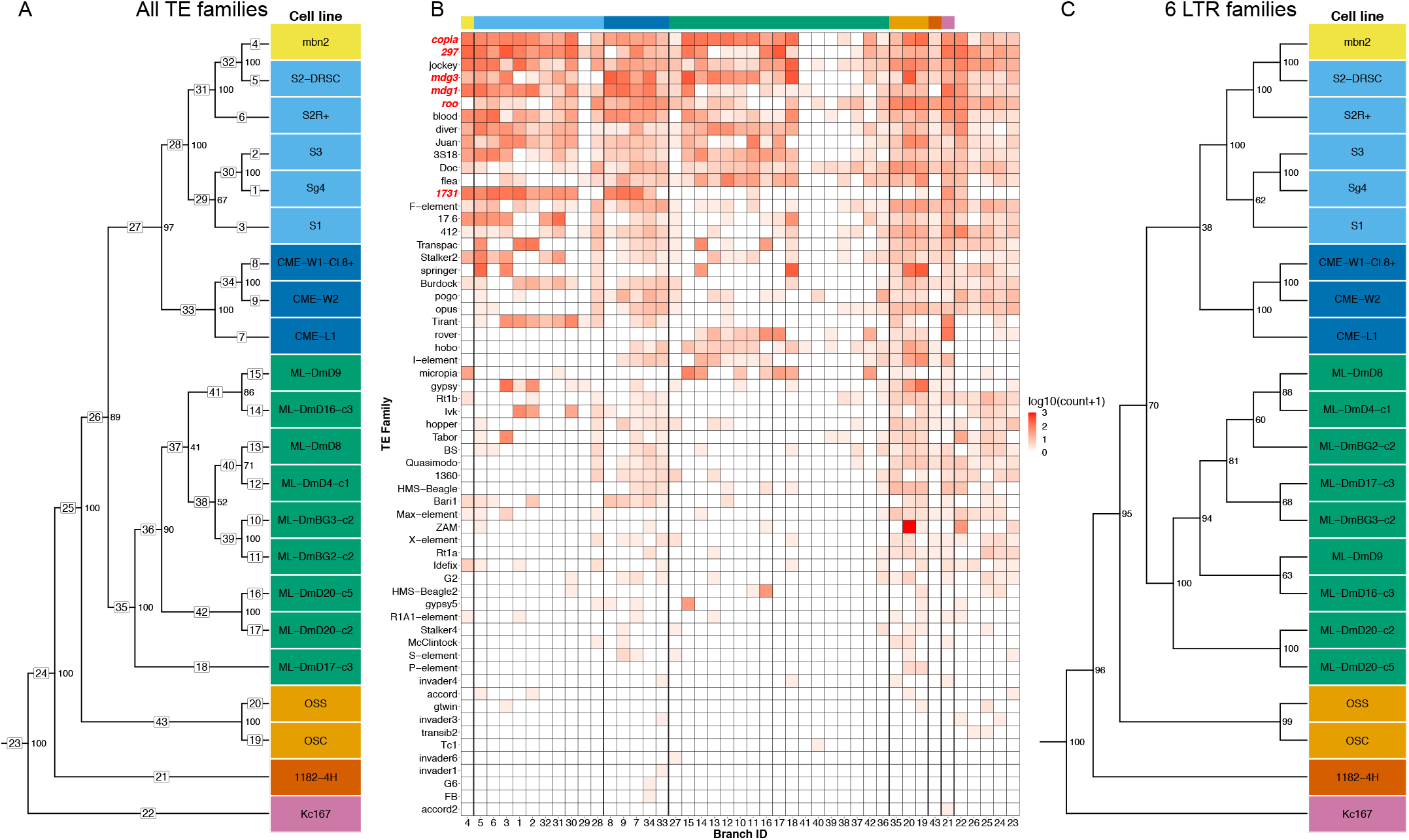
A small subset of LTR retrotransposon families can identify *Drosophila* cell lines. (A) Dollo parsimony tree of 22 *Drosophila* cell lines (without replicates) based on non-reference TE predictions for all 125 *D. melanogaster* TE families. Samples are colorized by lab origin as in Fig 2. Numbers inside boxes on branches indicate branch ID, and numbers beside nodes indicate percent support based on 100 bootstrap replicates. (B) Heatmap showing the number of non-reference TE insertion gain events per family on each branch of the tree in panel (A) based on ancestral state reconstruction using Dollo parsimony. The heatmap is colorized by log-transformed (log10(count+1)) number of gains per family per branch, sorted top to bottom by overall non-reference TE insertion gains per family across all branches, and sorted left to right into clades representing lab origin with lab origin clade color codes indicated at the top of the heatmap. The six diagnostic LTR retrotransposon families used in panel (C) are highlighted in red. (C) Dollo parsimony tree of 22 *Drosophila* cell lines (without replicates) based on non-reference predictions of six LTR retrotransposon families (297, copia, mdg3, mdg1, roo and 1731). Numbers beside nodes indicate percent support based on 100 bootstrap replicates.

Based on these results, we next evaluated whether a small, experimentally-tractable subset of TE families is sufficient to cluster and identify *Drosophila* cell lines. For this analysis, we focused on LTR retrotransposon families since this type of TE inserts with intact termini and therefore provide reliable 5’ and 3’ junctions for targeted PCR-based enrichment protocols (Smukowski Heil *et al.* 2021). We used the pattern of family- and branch-specific TE insertion to heuristically guide selection of a subset of six LTR retrotransposon families (copia, 297, mdg3, mdg1, roo, 1731; TE family names highlighted in red in Fig 3B), which defined unique TE profiles for each cell line and generated the same major patterns of *Drosophila* cell line clustering as the genome-wide dataset of all 125 TE families (3C). Finally, we tested whether a cell line sample (not used in the tree construction) can be accurately identified on the basis of its six-family TE profile. To do this, we used the six-family TE tree derived from the non-redundant set of primary replicates as a backbone to constrain Dollo parsimony searches including one additional “secondary” replicate for each of the 12 secondary replicates from the nine cell lines in the expanded dataset with secondary replicates. In 100% of cases (12/12), the additional secondary replicate clustered most closely with the primary replicate from the same cell line (Fig. S7). In 10/12 cases, the bootstrap support for the clustering of replicates was 100%, and the remaining two cases (both for CME-W1-Cl.8+) had lower bootstraps (≥64%) presumably because of the short read length for these secondary replicates (50bp). This proof-of-principle analysis indicates that TE insertions from a small subset of LTR retrotransposon families can accurately identify *Drosophila* cell line samples, and that only a subset of “diagnostic” TE families are needed to develop a targeted PCR-based enrichment protocol for *Drosophila* cell line authentication.

### TE profiles provide insight into Drosophila ovarian cell line history

The observation that different TE families are amplified in distinct *Drosophila* cell lines raises the question of whether a single TE family could diagnostically mark the identity of a *Drosophila* cell line or sub-line. One such candidate for this possibility is the retroviral-like LTR retrotransposon ZAM in the closely related OSS and OSC ovarian somatic cell lines (Niki *et al.* 2006; Saito *et al.* 2009). As shown above, we observed a massive increase in ZAM insertions in OSS cells relative to the OSC cell line (branches 19 and 20 in Fig 3A and B), supporting previous findings by Sytnikova *et al*. (2014). However, Sytnikova *et al*. (2014) also reported that ZAM amplification did not occur in all OSS sub-lines, only in a contemporary sub-line of OSS cells (called OSS_C), but not in a putatively early passage sub-line of OSS cells (called OSS_E).

To address whether ZAM proliferation is restricted to a subset of OSS sub-lines or is in fact a specific marker for all OSS sublines, we performed an integrated analysis of TE predictions in WGS data from six OSS and OSC samples from our and two previous studies (Sienski *et al.* 2012; Sytnikova *et al.* 2014). To formulate alternative hypotheses and guide interpretation of our results, we first compiled the reported provenance of these six OSS and OSC cell line samples. As shown in Fig. 4A, the ultimate ancestor of all OSS and OSC cell lines is a cell line composed of germline and somatic ovarian cell types called fGS/OSS (Niki *et al.* 2006). fGS/OSS cells were subsequently selected in the Niki lab to remove germline-marked stem cells to create the ancestor of the OSS (ovarian somatic sheet) cell line. The Niki lab sent two batches of OSS cells to the Lau lab in 2007 (Nelson Lau, personal communication): one was expanded and continuously cultured to become the OSS_C sub-line; the other was briefly cultured and stored as a cryopreserved culture for many years, then thawed and sequenced in 2013 creating the OSS_E sample (Sytnikova *et al.* 2014). Our sample of OSS cells comes from an independent sub-line donated by the Niki lab to the DGRC in 2010 (OSS_DGRC). The Niki lab also sent fGS/OSS cells to the Siomi lab, who independently selected against germline cells to create another somatic cell line called OSC (ovarian somatic cells) (Saito *et al.* 2009). OSC cells were sent by the Siomi lab in 2010 separately to the Lau (OSC_C) and Brennecke (OSC_E) labs, and were later donated by the Siomi lab to the DGRC in 2019 (OSC_DGRC).

**Figure 4.**
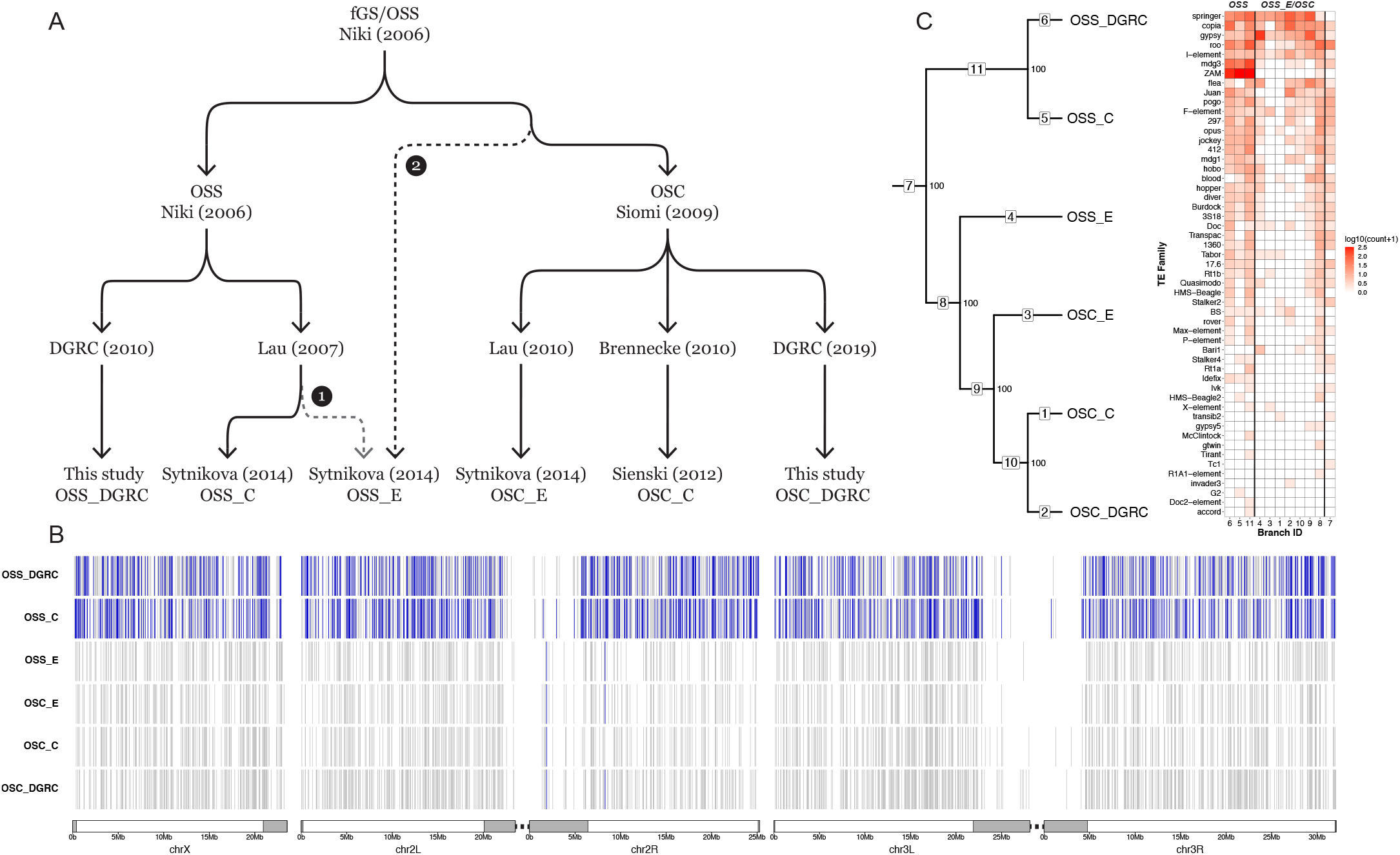
ZAM proliferation reveals OSS cell line identity. (A) Key events in the history of OSS and OSC cell line creation and distribution. Dotted lines represent alternative hypotheses for the identity of OSS_E. Branch 1 represents the reported provenance that hypothesizes OSS_E is an early diverging OSS sub-line; branch 2 hypothesizes that OSS_E approximates an ancestral state of the OSC cell line. (B) Genome-wide non-reference TE insertion data for six ovarian cell lines with ZAM insertions highlighted in blue and all other TE families in grey. (C) Dollo parsimony tree of ovarian cell lines based on all non-reference TE predictions. Numbers inside boxes on branches indicate branch ID, and numbers beside nodes indicate percent support based on 100 bootstrap replicates. (left). Heatmap showing the number of non-reference TE insertion gain events per family on each branch of the tree based on ancestral state reconstruction using Dollo parsimony. The heatmap is colorized by log-transformed (log10(count+1)) number of gains per family per branch, sorted top to bottom by overall non-reference TE insertion gains per family across all branches and sorted left to right into the *bona fide* OSS and OSS_E/OSC clusters (right).

Because WGS data from Sienski *et al*. (2012) and Sytnikova *et al*. (2014) is single-ended, integrated analysis of ovarian cell lines required a different TE prediction strategy than the one used for analysis of the paired-end datasets above. Preliminary analyses revealed that some single-end TE predictors (e.g. ngs_te_mapper, RelocaTE) (Linheiro and Bergman 2012; Robb *et al.* 2013) severely under-predicted insertions specifically for the ZAM family in the DGRC OSS sample relative to TEMP results based on paired-end data (Fig. S8). Additionally, our analysis of OSS and OSC samples ultimately required tracking intra-sample TE allele frequencies, which is not available in other TE predictors that use single-end data (e.g. TIDAL) (Rahman *et al.* 2015). Thus, we developed a new implementation of the single-end TE predictor originally described in in Linheiro and Bergman (2012) called ngs_te_mapper2 (https://github.com/bergmanlab/ngs_te_mapper2) that improves speed and sensitivity relative to the original version and has been extended to estimate intra-sample TE allele frequencies (Fig. S9; Table S4, Table S5; see Supplementary Text for details).

Using normalized datasets to optimize resolution of closely related sub-lines, we predicted non-reference TE insertions in all OSS and OSC sub-lines with ngs_te_mapper2 (File S4). These results revealed that ZAM has proliferated massively in the OSS_DGRC and OSS_C sub-lines (553 and 630 copies, respectively, in euchromatic regions), but is present in only one or two copies in OSS_E and all OSC sub-lines (Fig. 4B). The abundance of ZAM in these ovarian cell lines is more than 10-fold higher than fly strains where ZAM has been mobilized because of deletions in the *flamenco* piRNA locus (Leblanc *et al.* 1999; Zanni *et al.* 2013) or because of multigenerational knockdown of the piRNA effector protein *piwi* (Barckmann *et al.* 2018; Mohamed *et al.* 2020).

Under the “reported provenance” hypothesis that OSS_E and OSS_C share a more recent common ancestor than they do with OSS_DGRC (branch 1; Fig 4A), this pattern of ZAM abundance can only be explained by unlikely scenarios such as a massive loss of ZAM insertions on the branch leading to OSS_E, or independent parallel amplifications of ZAM on the OSS_C and OSS_DGRC sub-lines. An alternative hypothesis to explain the pattern of ZAM abundance is motivated by another observation made by Sytnikova *et al*. (2014): OSS_E shares more TE insertions in common with OSC sub-lines (OSC_E and OSC_C) than it does with a contemporary OSS sub-line (OSS_C). This pattern is not expected under the reported provenance hypothesis and suggests that OSS_E may in fact be an OSC-like lineage, rather than an early passage OSS sub-line. Under this alternative “uncertain provenance” hypothesis (branch 2; Fig 4A), the only *bona fide* OSS sub-lines would be OSS_C and OSS_DGRC, and ZAM proliferation could truly be a diagnostic marker of OSS cell line identity.

To test these alternative hypotheses, we used ngs_te_mapper2 predictions as input to cluster OSS and OSC sub-lines using Dollo parsimony. We found two highly supported clusters, one containing only the OSS_C plus OSS_DGRC sub-lines and the other containing OSS_E plus all OSC sub-lines (Fig. 4C, File S5). Ancestral state reconstruction clearly demonstrated that high ZAM abundance is restricted to the cluster containing OSS_C and OSS_DGRC sub-lines. The only two ZAM insertions that are found in OSS_E and OSC sub-lines are both shared by multiple sub-lines and therefore likely inserted in a common ancestor of the entire clade (Fig. 4B, File S6). We verified that the clustering relationships among OSS and OSC sub-lines were not solely driven by the ZAM amplification by repeating our clustering analysis excluding ZAM insertions, obtaining the same topology as in the complete dataset (Fig. S10A).

Further support for the hypothesis that OSS_E is an OSC-like lineage can be found in patterns of SNP and CNV variation in these cell line genomes (Fig. S10B and S10C). OSS_C and OSS_DGRC have essentially identical BAF profiles across the entire genome (Fig. S10B). In contrast, OSS_E and OSC sublines share a BAF profile everywhere but the distal regions on chromosome arms 2L, 3L and 3R (Fig. S10B, Fig. 5A). BAF profiles on all of chromosome X and arm 2R clearly differentiate OSS_C and OSS_DGRC (heterozygous) from OSS_E and OSC sub-lines (homozygous) (Fig. S10B). Likewise, CNV profiles support the clustering of OSS_C with OSS_DGRC and OSS_E with the OSC sub-lines. OSS_C and OSS_DGRC share a large deletion on chromosome X not found in OSS_E plus OSC sublines, and OSS_E plus the OSC sub-lines share a smaller deletion on chromosome arm 3L not found in OSS_C or OSS_DGRC (Fig. S10C). Based on these results, we conclude that OSS_E is a divergent lineage of OSC cells rather than early passage OSS cells, that ZAM amplification truly marks *bona fide* OSS cell lines (include the OSS line distributed by the DGRC), and that ngs_te_mapper2 TE predictions based on single-end WGS data can be effectively used to cluster *Drosophila* cell lines and reveal aspects of cell line history.

**Figure 5.**
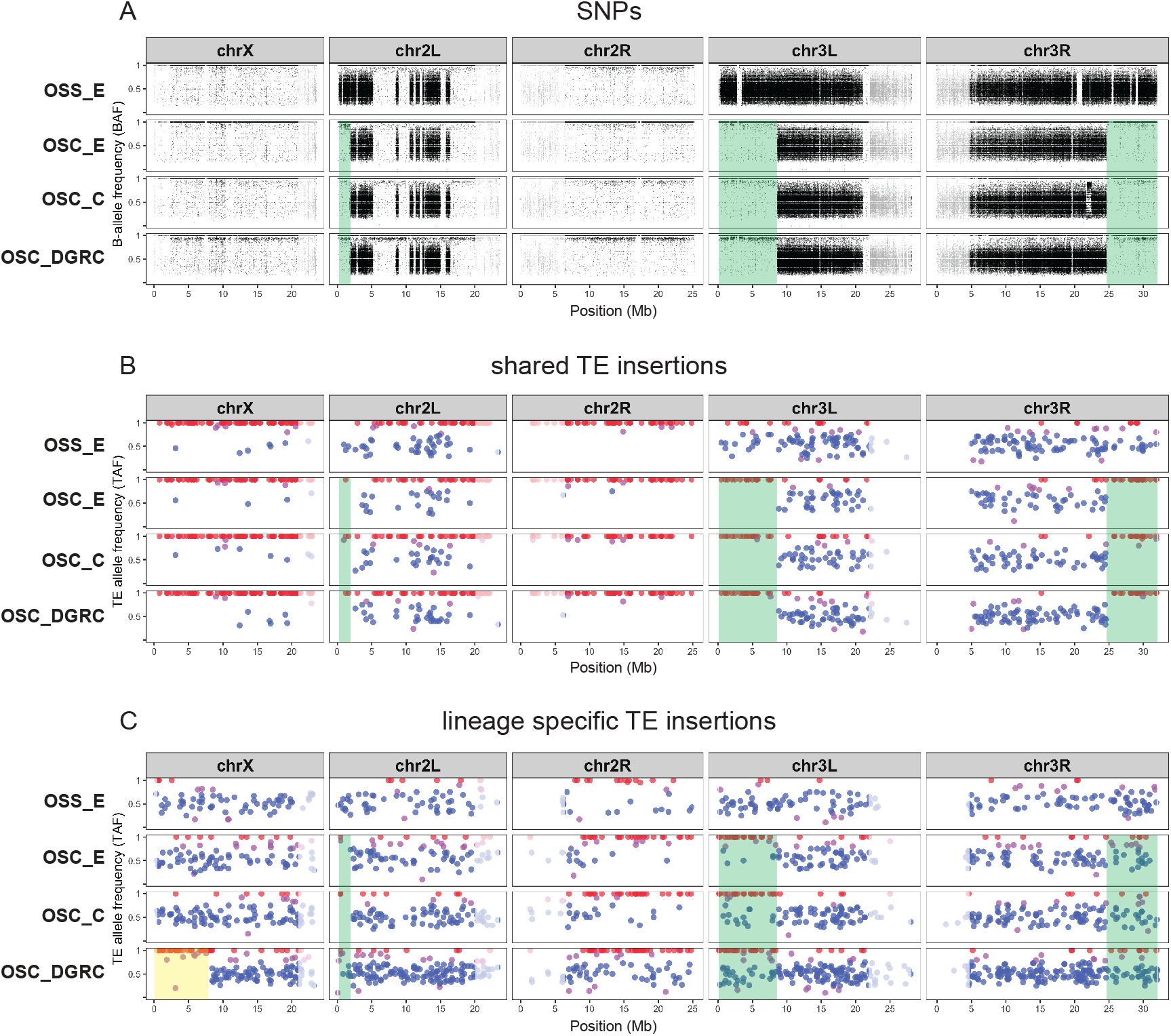
Loss of heterozygosity and ongoing transposition shape TE profiles in *Drosophila* ovarian somatic cell lines. Intrasample allele frequency profiles for OSS_E and OSC sub-lines based on (A) SNP variants, (B) TE insertions shared by OSS_E and OSC sub-lines, and (C) lineage specific TE insertions restricted to only OSS_E or the OSC sub-lines. SNPs and TE insertions in highly-repetitive low recombination regions are shaded in grey. For a given SNP, the B-allele frequency (BAF) was determined as the coverage of reads supporting non-reference allele divided by total coverage at that position. Regions of heterozygosity in a diploid genome are shown in BAF profiles where clusters of SNPs have allele frequencies centered around 0.5. Green shading indicates distal LOH regions defined by more extensive patterns of SNP heterozygosity in OSS_E relative to OSC sub-lines. TE insertions are classified as being homozygous (red), heterozygous (blue), or undefined (purple) based on allele frequencies estimated by ngs_te_mapper2. Yellow shading indicates LOH regions based on runs of homozygous TE insertions in OSC_DGRC relative to other OSC sub-lines.

### Loss of heterozygosity impacts TE profiles in Drosophila cell culture

Re-interpreting OSS_E as a divergent lineage of OSC cells requires explaining both the similarity and distinctness of its TE, BAF and CNV profiles from other OSC sub-lines. Two observations led us to hypothesize that OSS_E approximates an ancestral state of current OSC sub-lines. First, OSS_E occupies a basal position in the OSS_E plus OSC cluster based on TE profiles (Fig. 4C). Second, the BAF profile for OSS_E shows heterozygosity that extends in the distal regions of chromosome arms 2L, 3L and 3R relative to OSC sub-lines (green shading, Fig. 5A). We propose that differences in BAF profiles in these distal regions are caused by loss of heterozygosity (LOH) that occurred in an ancestor of all OSC sub-lines after divergence from the lineage leading to OSS_E. We infer that these distal LOH events were caused by mitotic recombination events rather than hemizygosity due to deletion, since copy number in distal LOH regions is the same in OSS_E and OSC sub-lines (Fig. S10C).

If this evolutionary scenario is correct, shared TEs (which inserted prior to the divergence of OSS_E and OSC sub-lines) that are heterozygous in OSS_E are predicted to be homozygous in OSC sub-lines in distal LOH regions, but should maintain heterozygosity elsewhere in the genome. To test these predictions, we used intra-sample allele frequency estimates from ngs_te_mapper2 to classify the zygosity of TE insertions shared by OSS_E and OSC sub-lines. Evaluation of our classifier on simulated genomes revealed it had *>*91% precision and crucially never falsely classified heterozygous insertions as homozygous (Table S6), and is thus conservative with respect to detection of LOH using TE insertions. As predicted under our model, we observed that there are many shared TE insertions in distal LOH regions that are heterozygous in OSS_E but virtually all TE insertions in these regions are homozygous in OSC sub-lines (green shading, Fig. 5B). Outside of distal LOH regions, shared TE insertions that are heterozygous in OSS_E generally retain heterozygosity in OSC sub-lines (Fig. 5B). In contrast, we observe that many lineage-specific TE insertions (which occurred after the divergence of OSS_E and OSC sub-lines) are heterozygous in OSC sub-lines in distal LOH regions (green shading, Fig. 5C). Together these results support the inferences that OSS_E approximates an ancestral state of current OSC sub-lines, that LOH events can cause fixation of previously heterozygous TE insertions in *Drosophila* cell lines, and that ongoing transposition in *Drosophila* cell culture can restore genetic variation in regions where previous large-scale LOH events have eliminated ancestral SNP or TE insertion variation.

Contrasting patterns of genetic variation between OSS_E and OSC sub-lines in distal regions of chromosome arms 2L, 3L and 3R provided the initial evidence for LOH due to mitotic recombination as mechanism of genome evolution in *Drosophila* cell culture. Assuming that the genome-wide heterozygosity observed in *bona fide* OSS sub-lines is ancestral (Fig. S10B), the lack of SNP heterozygosity on all of chromosome X and arm 2R in OSS_E and OSC sub-lines (Fig. 5A) supports the inference of additional whole-arm LOH events in the common ancestor of all of these sub-lines. Consistent with the prediction of whole-arm LOH in the ancestor of all OSS_E and OSC sub-lines followed by ongoing transposition in cell culture, we observe that most shared TE insertion on chromosome X and arm 2R are homozygous (Fig. 5B), while lineage-specific TE insertions are heterozygous (Fig. 5C). Intriguingly, and in contrast to other OSC sub-lines, we also observe that lineage-specific TE insertions on the distal eight megabases of chromosome X in OSC_DGRC are almost all homozygous (yellow shading, Fig. 5C). This observation can be explained by a secondary LOH event in the distal region of chromosome X that occurred recently only in the OSC_DGRC lineage. In this case, heterozygosity restored by ongoing TE insertion in *Drosophila* cell culture allows detection of a subsequent LOH events in the same genomic region that cannot be detected using SNP variation.

As LOH has not previously been reported as a mechanism of genome evolution in *Drosophila* cell culture, we sought to find additional evidence for this process by inspecting BAF profiles for other *Drosophila* cell lines in the expanded dataset. This led us to another potential case for LOH defined by SNPs on chromosome arms 2R and 3L of the CME-W2 and CME-W1-Cl.8+ cell lines (Fig. S3B, Fig. S11A). As with OSS_E, we propose that the more extensive heterozygous BAF profile on these chromosome arms in CME-W2 represents the pre-LOH ancestral-like state, and the homozygous BAF profile of CME-W1-Cl.8+ represents the post-LOH derived state. This scenario is consistent with the reported establishment of CME-W1-Cl.8+ from a single cloned cell of a polyclonal cell line (CME-W1) with the same ancestral genotype as CME-W2 (Currie *et al.* 1988; Peel and Milner 1990). The lack of difference in copy number profiles on chromosome arms 2R and 3L of CME-W2 and CME-W1-Cl.8+ suggests these events were also due to mitotic recombination (Fig. S3C). As predicted under the LOH model, we observed many TE insertions shared by CME-W2 and CME-W1-Cl.8+ are heterozygous in CME-W2 but are nearly all homozygous in CME-W1-Cl.8+ in LOH regions (Fig. S11B). Like in OSC sub-lines, we also observed many heterozygous TE insertions that are specific to CME-W1-Cl.8+ in LOH regions (Fig. S11C), consistent with recovery of TE insertion variation after LOH. Evidence for LOH in distinct cell lines developed in two different labs generalizes the inference that LOH shapes TE profiles in *Drosophila* cell lines, and suggests that LOH as a mechanism of genome evolution in *Drosophila* culture is not dependent on the genetic background of ancestral fly donor.

## Conclusions

Here we demonstrate that TE insertion profiles can successfully identify *Drosophila* cell lines and use this finding to clarify several aspects of cell line provenance in *Drosophila*. The success of this approach validates our basic model for how the joint processes of germline transposition in whole flies and somatic transposition in cell culture create TE profiles that uniquely mark *Drosophila* cell lines (Fig. 1). We also show that TE insertion profiles can shed light on the evolutionary history of *Drosophila* cell lines derived from a common ancestral cell line, and that LOH due to mitotic recombination is an additional mechanism of genome evolution in cell culture that adds complexity to our basic model (Fig. 6). During cell culture, mitotic recombination events purge ancestral variation distal to cross-over breakpoints, causing previously heterozygous SNPs and TE insertions to become fixed or lost within a cell line genome (green shading). Ongoing transposition in cell culture after LOH leads to the relatively rapid recovery of TE but not SNP heterozygosity, allowing secondary LOH events to be identified using TE insertions in regions that have previous lost ancestral variation due to primary LOH events (yellow shading). The emerging model of TE evolution in cell culture motivated by results presented here has direct implications for the development of protocols for cell line identification in *Drosophila* and contributes to our general understanding of the mechanisms of genome evolution in cell lines derived from multicellular organisms.

**Figure 6.**
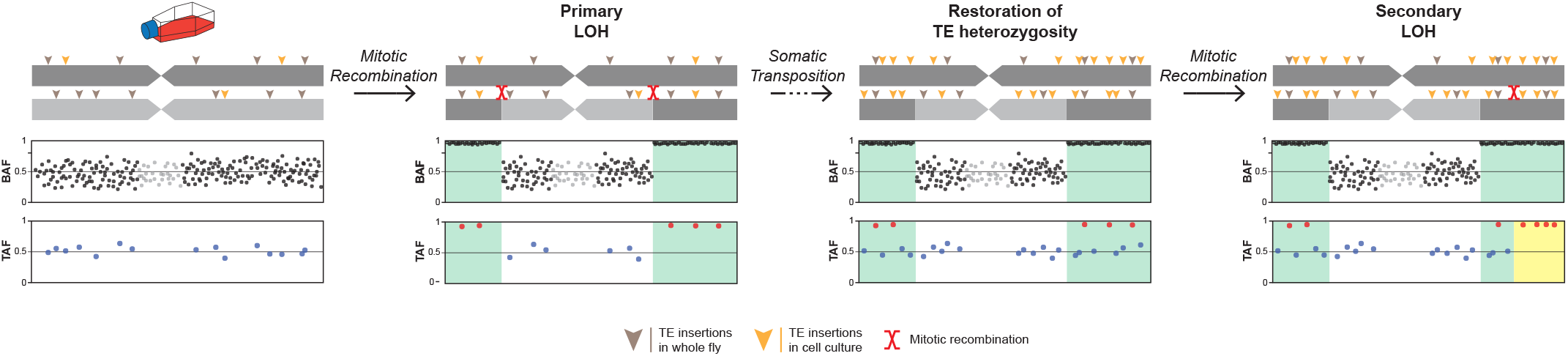
Schematic model of how loss of heterozygosity and somatic transposition interact to shape TE profiles in *Drosophila* cell line genomes. Mitotic recombination in cell culture between homologous chromosomes causes LOH of pre-existing heterozygous SNP and TE variants, revealed respectively by B-allele frequency (BAF) and TE-allele frequency (TAF) profiles, in regions distal to cross-over breakpoints (green shading). Ongoing transposition in cell culture leads to accumulation of new haplotype-specific heterozygous TE insertions inside and outside of primary LOH regions. Restoration of TE heterozygosity allows detection of secondary LOH events (yellow shading) in regions of the genome that have previously undergone primary LOH events. The model depicts a simplified case of diploidy, when in reality cell culture genomes can have complex genome structure due to polyploidy and segmental aneuploidy.

## Materials and Methods

### Genome sequencing

Public genome sequencing data for 26 samples of 18 *Drosophila* cell lines were obtained from the modENCODE project (Lee *et al.* 2014). Frozen stocks of eight additional samples from six *Drosophila* cell lines (mbn2, Sg4, ML-DmBG3-c2, ML-DmBG2-c2, OSS and OSC) were obtained from the *Drosophila* Genomics Resource Center (DGRC), the Gorski lab (Canada’s Michael Smith Genome Sciences Centre, BC Cancer) and the Strand lab (University of Georgia). DNA extractions were performed using Qiagen Blood and Tissue kit (Cat# 69504) for the mbn2 sample from the Strand lab and using the Zymo-Quick kit (Cat#D4068) for all other samples. Purified DNA was analyzed by Qubit and Fragment Analyzer to determine the concentration and size distribution, respectively. Samples were normalized to the same concentration before preparing libraries with the KAPA Hyper Prep Kit (Cat# KK8504). During library prep, DNA was fragmented by acoustic shearing with Covaris E220 Evolution before end repair and A-tailing. Single indices were ligated to DNA fragments. Libraries were purified and cleaned with Solid Phase Reversible Immobilization (SPRI) beads before PCR amplification. Final libraries underwent an additional round of bead cleanup before being assessed by Qubit, qPCR (KAPA Library Quantification Kit Cat# KK4854), and Fragment Analyzer. Libraries were then sequenced in paired-end 150bp mode on an Illumina NextSeq500 high output flowcell and demultiplexed using bcl2fastq. Metadata, sequencing statistics, and SRA accession numbers for all cell line DNA-seq samples used in this study can be found in Table S1.

### Detection of non-reference TE insertions using paired-end sequencing data

Paired-end sequencing data from the modENCODE project (Lee *et al*. 2014) and our study was used as input to seven methods designed to detect non-reference TE insertions in *Drosophila* (Linheiro and Bergman 2012; Kofler *et al*. 2012; Zhuang *et al*. 2014; Kofler *et al*. 2016; Adrion *et al*. 2017; Yu *et al*. 2021) using McClintock (revision 40863acf11052b18afb4cdcd7b1124de48cba397; options: -m “trimgalore, popoolationte, popoolationte2, temp, temp2, teflon, ngs_te_mapper, ngs_te_mapper2”) (Nelson *et al.* 2017). Additionally, we predicted non-reference TE insertions using a version of TIDAL 1.2 (Rahman *et al.* 2015; Yang *et al.* 2021) that was modified to output results in a format compatible with results from McClintock (https://github.com/pbasting/TIDAL1.2, revision 2d110b17b3b287dbc1ceb67c87fe171d15095c84). The reference genome for these analyses was comprised of the major chromosome arms from the *D. melanogaster* dm6 assembly (chr2L, chr2R, chr3L, chr3R, chr4, chrM, chrY, and chrX) and the TE library was the Berkeley *Drosophila* Genome Project canonical TE dataset v10.1 (https://github.com/bergmanlab/transposons/blob/master/releases/D_mel_transposon_sequence_set_v10.1.fa; revision f94d53ea10b95c9da99258ac2336ce18871768e9).

Paired-end samples analyzed here vary substantially in read length (50-151 bp) and depth of coverage (5X-136X) (Table S1). We chose not to normalize input datasets by downsampling to the lowest read length and coverage to avoid reducing sensitivity of non-reference TE detection methods for higher quality samples. Using complete samples allowed us to observe that the number of non-reference TE predictions per sample (Table S2) showed a strong dependence on read length (Fig. S1) or coverage (Fig. S2) for all methods besides TEMP (Zhuang *et al.* 2014). Thus, we used TEMP predictions with default McClintock filtering (retain only 1p1 predictions with >0.1 intra-sample allele frequency cutoff) for the global analysis of the modENCODE-only and expanded (modENCODE plus new samples) datasets. To resolve details of the relationship among mbn2 sub-lines, we used read length and coverage normalized mbn2 samples with relaxed filtering criteria for TEMP predictions (retain all 1p1/2p/singlton predictions with no intra-sample allele frequency cutoff).

### Detection of non-reference TE insertions using single-end sequencing data

Single-end sequencing data for OSS and OSC cell line samples from two previous studies (Sienski *et al.* 2012; Sytnikova *et al*. 2014) and forward reads from our paired-end samples were used to predict non-reference TE insertions using ngs_te_mapper2 (https://github.com/bergmanlab/ngs_te_mapper2) in McClintock (revision 40863acf11052b18afb4cdcd7b1124de48cba397; options:-m “trimgalore, coverage, ngs_te_mapper2, map_reads”) (Nelson *et al.* 2017). ngs_te_mapper2 is a re-implementation of the non-reference TE detection method initially reported in Linheiro and Bergman (2012) that improves speed and sensitivity and has been extended to estimate TE allele frequency (see Supplementary Text for details). Reference genome and TE library files used for McClintock runs on single-end sequencing data were the same as used above for paired-end sequencing data. Because ngs_te_mapper2 detection rates and allele frequency estimates are sensitive to read length and depth of coverage (see Supplementary Text), reads from single-end sequencing data and the forward read of our paired-end sequencing data were normalized by trimming all reads to 100bp using fastp v0.20.1 (Chen *et al.* 2018) and downsampling to the lowest coverage sample (14X) using seqtk v1.3 (Li 2015).

### Classification of intra-sample TE insertion allele frequency

To predict whether TE insertions within OSS and OSC cell line samples were heterozyogous or homozygous, we built a classifier that uses allele frequencies estimated by ngs_te_mapper2 from single-end sequencing data as input. A non-reference TE insertion was predicted to be heterozyogous if the intra-sample allele frequency estimated by ngs_te_mapper2 is between 0.25 to 0.75 and predicted to be homozygous if the intra-sample allele frequency is greater than or equal to 0.95. TE insertions with intra-sample allele freqeuncies outside these ranges were considered unclassified. The classifier was benchmarked using synthetic homozygous and heterozygous WGS datasets created with wgsim v0.3.1-r13 using the ISO1 (dm6) and A4 (GCA_003401745.1) (Chakraborty *et al.* 2018) genome assemblies as input. The classifier yields >91% precision using input from the results of ngs_te_mapper2 applied to the simulated datasets (see Supplementary Text for details).

### Identification of orthologous TE insertions

Because positional resolution of non-reference TE predictions is inexact (Nelson *et al*. 2017), we identified a high-quality set of orthologous non-reference TE insertion loci as follows. Genome-wide non-redundant BED files of non-reference TE predictions generated by McClintock were filtered to exclude TEs in low recombination regions using boundaries defined by Cridland *et al*. (2013) lifted over to dm6 coordinates. Normal recombination regions included in our analyses were defined as chrX:405967–20928973, chr2L:200000–20100000, chr2R:6412495– 25112477, chr3L:100000–21906900, chr3R:4774278–31974278. We restricted our analysis to normal recombination regions, since low recombination regions have high reference TE content which reduces the ability to predict non-reference TE insertions (Bergman *et al.* 2006; Manee *et al.* 2018). We also excluded INE-1 family from our analysis, as this family is reported to be inactive for millions of years (Singh and Petrov 2004; Wang *et al.* 2007). Non-reference TE predictions in high recombination from all samples were then clustered into orthologous loci using BEDtools cluster v2.26.0 enforcing predictions within each cluster to be on the same strand (option -s) (Quinlan and Hall 2010). Orthologous loci were then filtered using the following criteria: 1) retain only a single TE family per locus; 2) retain only a single TE prediction per sample per locus; and 3) retain TE predictions only from long-terminal repeat (LTR) retrotransposon, LINE-like retrotransposon or DNA transposon families. For clustering of paired-end samples, we imposed the additional filtering requirement that all clusters include at least sample per locus with a TEMP 1p1 prediction.

### Clustering and identification of cell line samples using TE insertion profiles

Non-reference TE predictions at orthologous loci were then converted to a binary presence/absence matrix in order to cluster cell lines on the basis of their TE insertion profiles. Cell line clustering was performed using Dollo parsimony in PAUP (v4.0a168) (Swofford 2003). Dollo parsimony analyses were conducted using heuristic searches with 50 replicates. A hypothetical ancestor carrying the assumed ancestral state for each locus (absence) was included as a root in the analysis (Batzer and Deininger 2002). “DescribeTrees chgList=yes” option was used to assign character state changes to branches in the tree. Node support for the most parsimonious tree was evaluated by integrating 100 bootstrap replicates generated by PAUP using SumTrees (Sukumaran and Holder 2010).

Identification of a cell line sample was performed by adding its TE profile to a binary presence/absence matrix of “primary replicates” of 22 non-redundant *Drosophila* cell line samples and performing cell line clustering using the same approach mentioned above. A phylogenetic tree of the 22 non-redundant primary *Drosophila* cell line samples was used as a backbone topological constraint during a heuristic searches for the most parsimonious tree that included one additional “secondary replicate”. Node support for the most parsimonious tree was evaluated by integrating 100 bootstrap replicates without topological constraints.

### B-allele frequency and copy number analysis

BAM files generated by McClintock were used for variant calling using bcftools v1.9 (Li 2011). Indels were excluded from variant calling, leaving only single-nucleotide polymorphisms (SNPs) in the VCF file. For a given SNP, the B-allele frequency (BAF) was determined as the coverage of reads supporting non-reference allele divided by total coverage at that position using the DP4 field.

BAM files generated by McClintock were also used to generate copy number variant (CNV) profiles for non-overlapping 10kb windows of the dm6 genome using Control-FREEC (v11.6) (Boeva *et al.* 2012). Windows with less than 85% mappability were excluded from the analysis based on mappability tracks generated by GEM (v1.315 beta) (Derrien *et al.* 2012). The baseline ploidy was determined by normalized DNA read density of 10 kb windows following Lee *et al*. (2014). The sex information was determined from relative read density between chromosome X and autosomes. The minimum and maximum expected value of the GC content was set to be 0.3 and 0.45, respectively.

### Clustering of cell line samples based on transcriptomes

Total RNA sequencing samples for 17 *Drosophila* cell lines with 100bp paired-end reads were obtained from (Stoiber *et al.* 2016) and from the modENCODE *D. melanogaster* transcriptome sequencing project (Brown *et al.* 2014). SRA accession numbers for all cell line RNA-seq samples used in this analysis can be found in (Table S3). Transcript abundances for protein-coding genes were quantified in unit of transcripts per million (TPM) using kallisto quant v0.46.2 (Bray *et al.* 2016) using the release 6.32 version of the *D. melanogaster* transcript coding sequences corresponding to Ensembl genes from Ensembl release 103 (http://ftp.ensembl.org/pub/release-103/fasta/drosophila_melanogaster/cds/Drosophila_melanogaster.BDGP6.32.cds.all.fa.gz) (Yates *et al*. 2020). Transcript-level abundance estimates were summarized into gene-level abundance estimates using the release 6.32 version of the *D. melanogaster* gene annotation from Ensembl release 103 (http://ftp.ensembl.org/pub/release-103/gtf/drosophila_melanogaster/Drosophila_melanogaster.BDGP6.32.103.gtf.gz) using tximport v1.18.0 (Soneson *et al*. 2015). The summarized gene-level abundance matrix was log transformed and visualized using the Rtsne package v0.15 (Krijthe 2015).

## Data Availability

File S1 contains nonredundant bed files from McClintock runs using TEMP module on the expanded dataset including 34 *Drosophila* cell line samples. File S2 contains clustered TE profiles in the format of binary presence/absence data matrix including 34 *Drosophila* cell line samples. File S3 includes data matrix of the number of non-reference TE insertion gain events per family on each branch of the most parsimonious tree used for the heatmap in Fig. 3B. File S4 includes nonredundant bed files from McClintock runs using ngs_te_mapper2 module on the normalized OSS and OSC dataset. File S5 includes clustered TE profiles in the format of binary presence/absence data matrix including 6 OSS and OSC cell line samples. File S6 includes data matrix of the number of non-reference TE insertion gain events per family on each branch of the most parsimonious tree used for the heatmap in Fig. 4C. Raw sequencing data generated in our study is available in the SRA under BioProject PRJNA689777.

## Acknowledgements

We thank Nancy Go and Sharon Gorski (Canada’s Michael Smith Genome Sciences Centre, BC Cancer) and Michael Strand (University of Georgia) for supplying samples of mbn2 cells; Noah Workman and Magdy Alabady at the University of Georgia Genomics and Bioinformatics Core for assistance with Illumina library preparation and sequencing; Dan Hultmark (Umeå University), Julius Brennecke (Institute of Molecular Biotechnology, Vienna) and Nelson Lau (Boston University) for information about the history of mbn2, OSS and OSC cell lines; and Nelson Lau for helpful comments on the manuscript. This work was supported in part by resources and technical expertise from the Georgia Advanced Computing Resource Center, and by funding from a University of Georgia Research Education Award Traineeship (P.J.B.), the University of Georgia Research Foundation (C.M.B.), and NIH grant 2P40OD010949 (A.Z.).

## Literature Cited

Adrion, J. R., M. J. Song, D. R. Schrider, M. W. Hahn, and S. Schaack, 2017 Genome-wide estimates of transposable element insertion and deletion rates in Drosophila melanogaster. Genome Biol Evol 9: 1329–1340.

Almeida, J. L., K. D. Cole, and A. L. Plant, 2016 Standards for cell line authentication and beyond. PLOS Biology 14: e1002476. American Type Culture Collection Standards Development Organization Workgroup ASN-0002, 2010 Cell line misidentification: the beginning of the end. Nat Rev Cancer 10: 441–448.

Babic, Z., A. Capes-Davis, M. E. Martone, A. Bairoch, I. B. Ozyurt, et al., 2019 Incidences of problematic cell lines are lower in papers that use RRIDs to identify cell lines. eLife 8: e41676.

Barallon, R., S. R. Bauer, J. Butler, A. Capes-Davis, W. G. Dirks, et al., 2010 Recommendation of short tandem repeat profiling for authenticating human cell lines, stem cells, and tissues. In Vitro Cell Dev Biol Anim 46: 727–732.

Barckmann, B., M. El-Barouk, A. Pélisson, B. Mugat, B. Li, et al., 2018 The somatic piRNA pathway controls germline transposition over generations. Nucleic Acids Res 46: 9524–9536.

Batzer, M. A. and P. L. Deininger, 2002 Alu repeats and human genomic diversity. Nature Reviews Genetics 3: 370–379.

Bergman, C. M., H. Quesneville, D. Anxolabehere, and M. Ashburner, 2006 Recurrent insertion and duplication generate net-works of transposable element sequences in the Drosophila melanogaster genome. Genome Biol 7: R112.

Boeva, V., T. Popova, K. Bleakley, P. Chiche, J. Cappo, et al., 2012 Control-FREEC: a tool for assessing copy number and allelic content using next-generation sequencing data. Bioinformatics 28: 423–425.

Bray, N. L., H. Pimentel, P. Melsted, and L. Pachter, 2016 Nearoptimal probabilistic RNA-seq quantification. Nature Biotechnology 34: 525–527.

Brown, J. B., N. Boley, R. Eisman, G. E. May, M. H. Stoiber, et al., 2014 Diversity and dynamics of the Drosophila transcriptome. Nature 512: 393–399.

Capes-Davis, A., G. Theodosopoulos, I. Atkin, H. G. Drexler, A. Kohara, et al., 2010 Check your cultures! A list of cross-contaminated or misidentified cell lines. Int J Cancer 127: 1–8.

Castro, F., W. G. Dirks, S. Fähnrich, A. Hotz-Wagenblatt, M. Pawlita, et al., 2013 High-throughput SNP-based authentication of human cell lines. Int J Cancer 132: 308–314.

Chakraborty, M., N. W. VanKuren, R. Zhao, X. Zhang, S. Kalsow, et al., 2018 Hidden genetic variation shapes the structure of functional elements in Drosophila. Nat Genet 50: 20–25.

Charlesworth, B. and C. H. Langley, 1989 The population genetics of Drosophila transposable elements. Annu Rev Genet 23: 251–87.

Chen, S., Y. Zhou, Y. Chen, and J. Gu, 2018 fastp: an ultra-fast all-in-one FASTQ preprocessor. Bioinformatics 34: i884.–i890.

Cherbas, L., A. Willingham, D. Zhang, L. Yang, Y. Zou, et al., 2011 The transcriptional diversity of 25 Drosophila cell lines. Genome Res 21: 301–314.

Cridland, J. M., S. J. Macdonald, A. D. Long, and K. R. Thornton, 2013 Abundance and distribution of transposable elements in two Drosophila QTL mapping resources. Mol Biol Evol 30: 2311–2327.

Currie, D. A., M. J. Milner, and C. W. Evans, 1988 The growth and differentiation in vitro of leg and wing imaginal disc cells from Drosophila melanogaster. Development 102: 805–814.

Defendi, V., R. E. Billingham, W. K. Silvers, and P. Moorhead, 1960 Immunological and karyological criteria for identification of cell lines. JNCI: Journal of the National Cancer Institute 25: 359–385.

Derrien, T., J. Estelle, S. M. Sola, D. G. Knowles, E. Raineri, et al., 2012 Fast Computation and Applications of Genome Mappability. PLOS ONE 7: e30377.

Echalier, G., 1997 Drosophila Cells in Culture. Academic Press, San Diego, Calif. Echalier, G. and A. Ohanessian, 1969 Isolement, en cultures in vitro, de lignees cellulaires diploides de Drosophila melanogaster. Comptes rendus hebdomadaires des seances de l’Academie des Sciences 268: 1771–1773.

Gartler, S. M., 1967 Genetic markers as tracers in cell culture. Natl Cancer Inst Monogr 26: 167–195.

Gateff, E., 1977 [New mutants report.]. Drosophila Information Service 52: 4–5.

Gateff, E., L. Gissmann, R. Shrestha, N. Plus, H. Pfister, et al., 1980 Characterization of two tumorous blood cell lines of Drosophila melanogaster and the viruses they contain. Invertebrate Systems in Vitro Fifth International Conference on Invertebrate Tissue Culture, Rigi-Kaltbad, Switzerland, 1979 pp. 517–533.

Gilbert, D. A., Y. A. Reid, M. H. Gail, D. Pee, C. White, et al., 1990 Application of DNA fingerprints for cell-line individualization. Am J Hum Genet 47: 499–514.

Horbach, S. P. J. M. and W. Halffman, 2017 The ghosts of HeLa: How cell line misidentification contaminates the scientific literature. PLOS ONE 12: e0186281.

Huang, Y., Y. Liu, C. Zheng, and C. Shen, 2017 Investigation of cross-contamination and misidentification of 278 widely used tumor cell lines. PLOS One 12: e0170384.

Ilyin, Y. V., V. G. Chmeliauskaite, E. V. Ananiev, and G. P. Georgiev, 1980 Isolation and characterization of a new family of mobile dispersed genetic elements, mdg3, in Drosophila melanogaster. Chromosoma 81: 27–53.

Kofler, R., A. J. Betancourt, and C. Schlotterer, 2012 Sequencing of pooled DNA samples (pool-seq) uncovers complex dynamics of transposable element insertions in Drosophila melanogaster. PLOS Genet 8: e1002487.

Kofler, R., D. Gomez-Sanchez, and C. Schlotterer, 2016 PoPoolationTE2: comparative population genomics of transposable elements using pool-seq. Mol Biol Evol 33: 2759–2764.

Krijthe, J. H., 2015 Rtsne: T-Distributed Stochastic Neighbor Embedding using Barnes-Hut Implementation.

Larkin, A., S. J. Marygold, G. Antonazzo, H. Attrill, G. dos Santos, et al., 2021 FlyBase: updates to the Drosophila melanogaster knowledge base. Nucleic Acids Res 49: D899– D907.

Leblanc, P., B. Dastugue, and C. Vaury, 1999 The Integration Machinery of ZAM, a Retroelement from Drosophila melanogaster, Acts as a Sequence-Specific Endonuclease. J Virol 73: 7061–7064.

Lee, H., C. J. McManus, D.-Y. Cho, M. Eaton, F. Renda, et al., 2014 DNA copy number evolution in Drosophila cell lines. Genome Biol 15: R70.

Li, H., 2011 A statistical framework for SNP calling, mutation discovery, association mapping and population genetical parameter estimation from sequencing data. Bioinformatics 27: 2987–2993.

Li, H., 2015 seqtk.

Liang-Chu, M. M. Y., M. Yu, P. M. Haverty, J. Koeman, J. Ziegle, et al., 2015 Human biosample authentication using the high-throughput, cost-effective snptrace system. PLOS ONE 10: e0116218.

Linheiro, R. S. and C. M. Bergman, 2012 Whole genome resequencing reveals natural target site preferences of transposable elements in Drosophila melanogaster. PLOS One 7: e30008.

Lorsch, J. R., F. S. Collins, and J. Lippincott-Schwartz, 2014 Fixing problems with cell lines. Science 346: 1452–1453.

Luhur, A., K. M. Klueg, and A. C. Zelhof, 2019 Generating and working with Drosophila cell cultures: Current challenges and opportunities. Wiley Interdiscip Rev Dev Biol 8: e339.

Maaten, L. v. d., 2014 Accelerating t-SNE using tree-based algorithms. Journal of Machine Learning Research 15: 3221–3245.

Maaten, L. v. d. and G. Hinton, 2008 Visualizing data using t-SNE. Journal of Machine Learning Research 9: 2579–2605.

MacLeod, R. A., W. G. Dirks, Y. Matsuo, M. Kaufmann, H. Milch, et al., 1999 Widespread intraspecies cross-contamination of human tumor cell lines arising at source. Int J Cancer 83: 555– 563.

Manee, M. M., J. Jackson, and C. M. Bergman, 2018 Conserved noncoding elements influence the transposable element landscape in Drosophila. Genome Biol Evol 10: 1533–1545.

Masters, J. R., J. A. Thomson, B. Daly-Burns, Y. A. Reid, W. G. Dirks, et al., 2001 Short tandem repeat profiling provides an international reference standard for human cell lines. Proc Natl Acad Sci USA 98: 8012–8017.

Mohamed, M., N. T.-M. Dang, Y. Ogyama, N. Burlet, B. Mugat, et al., 2020 A Transposon Story: From TE Content to TE Dynamic Invasion of Drosophila Genomes Using the Single-Molecule Sequencing Technology from Oxford Nanopore. Cells 9: 1776.

Mohammad, T. A., Y. S. Tsai, S. Ameer, H.-I. H. Chen, Y.-C. Chiu, et al., 2019 CeL-ID: cell line identification using RNA-seq data. BMC Genomics 20: 81.

Morales, V., T. Straub, M. F. Neumann, G. Mengus, A. Akhtar, et al., 2004 Functional integration of the histone acetyltransferase MOF into the dosage compensation complex. The EMBO Journal 23: 2258–2268.

Nelson, M. G., R. S. Linheiro, and C. M. Bergman, 2017 Mc-Clintock: an integrated pipeline for detecting transposable element insertions in whole-genome shotgun sequencing data. G3 7: 2749–2762.

Nelson-Rees, W. A., D. W. Daniels, and R. R. Flandermeyer, 1981 Cross-contamination of cells in culture. Science 212: 446–452.

Niki, Y., T. Yamaguchi, and A. P. Mahowald, 2006 Establishment of stable cell lines of Drosophila germ-line stem cells. Proc Natl Acad Sci USA 103: 16325–16330.

O’Brien, S. U., G. Kleiner, R. Olson, and J. E. Shannon, 1977 Enzyme polymorphisms as genetic signatures in human cell cultures. Science 195: 1345–1348.

Parson, W., R. Kirchebner, R. Mühlmann, K. Renner, A. Kofler, et al., 2005 Cancer cell line identification by short tandem repeat profiling: power and limitations. FASEB J 19: 434–436.

Peel, D. J. and M. J. Milner, 1990 The diversity of cell morphology in cloned cell lines derived from Drosophila imaginal discs. Roux’s Arch Dev Biol 198: 479–482.

Potter, S. S., W. J. Brorein, P. Dunsmuir, and G. M. Rubin, 1979 Transposition of elements of the 412, copia and 297 dispersed repeated gene families in Drosophila. Cell 17: 415–427.

Quinlan, A. R. and I. M. Hall, 2010 BEDTools: a flexible suite of utilities for comparing genomic features. Bioinformatics 26: 841–842.

Rahman, R., G.-w. Chirn, A. Kanodia, Y. A. Sytnikova, B. Brembs, et al., 2015 Unique transposon landscapes are pervasive across Drosophila melanogaster genomes. Nucleic Acids Res 43: 10655–10672.

Ray, D. A., J. Xing, A.-H. Salem, and M. A. Batzer, 2006 SINEs of a nearly perfect character. Syst Biol 55: 928–935.

Ress, C., M. Holtmann, U. Maas, J. Sofsky, and A. Dorn, 2000 20-Hydroxyecdysone-induced differentiation and bapoptosis in the Drosophila cell line, l(2)mbn. Tissue and Cell 32: 464–477.

Rishishwar, L., L. Marino-Ramirez, and I. K. Jordan, 2017 Bench-marking computational tools for polymorphic transposable element detection. Brief Bioinformatics 18: 908–918.

Robb, S. M. C., L. Lu, E. Valencia, J. M. Burnette, Y. Okumoto, et al., 2013 The use of RelocaTE and unassembled short reads to produce high-resolution snapshots of transposable element generated diversity in rice. G3 3: 949–957.

Roy, S., J. Ernst, P. V. Kharchenko, P. Kheradpour, N. Negre, et al., 2010 Identification of functional elements and regulatory circuits by Drosophila modENCODE. Science 330: 1787–1797.

Saito, K., S. Inagaki, T. Mituyama, Y. Kawamura, Y. Ono, et al., 2009 A regulatory circuit for piwi by the large Maf gene traffic jam in Drosophila. Nature 461: 1296–1299.

Samakovlis, C., B. Asling, H. G. Boman, E. Gateff, and D. Hultmark, 1992 In vitro induction of cecropin genes — an immune response in a Drosophila blood cell line. Biochemical and Biophysical Research Communications 188: 1169–1175.

Schneider, I., 1972 Cell lines derived from late embryonic stages of Drosophila melanogaster. J Embryol Exp Morphol 27: 353– 365.

Schug, M. D., T. F. Mackay, and C. F. Aquadro, 1997 Low mutation rates of microsatellite loci in Drosophila melanogaster. Nat Genet 15: 99–102.

Schwartz, Y. B., T. G. Kahn, D. A. Nix, X.-Y. Li, R. Bourgon, et al., 2006 Genome-wide analysis of Polycomb targets in Drosophila melanogaster. Nature Genetics 38: 700–705.

Sienski, G., D. Dönertas, and J. Brennecke, 2012 Transcriptional silencing of transposons by Piwi and maelstrom and its impact on chromatin state and gene expression. Cell 151: 964–980.

Singh, N. D. and D. A. Petrov, 2004 Rapid sequence turnover at an intergenic locus in Drosophila. Molecular Biology and Evolution 21: 670–80.

Smukowski Heil, C., K. Patterson, A. Shang-Mei Hickey, E. Alcantara, and M. J. Dunham, 2021 Transposable element mobilization in interspecific yeast hybrids. Genome Biol Evol 13: evab033.

Soneson, C., M. I. Love, and M. D. Robinson, 2015 Differential analyses for RNA-seq: transcript-level estimates improve gene-level inferences. F1000Res 4: 1521.

Stanley, C. E. and R. J. Kulathinal, 2016 Genomic signatures of domestication on neurogenetic genes in Drosophila melanogaster. BMC Evolutionary Biology 16: 6.

Stoiber, M., S. Celniker, L. Cherbas, B. Brown, and P. Cherbas, 2016 Diverse Hormone Response Networks in 41 Independent Drosophila Cell Lines. G3 6: 683–694.

Sukumaran, J. and M. T. Holder, 2010 DendroPy: a Python library for phylogenetic computing. Bioinformatics 26: 1569– 1571.

Swofford, D., 2003 PAUP*: phylogenetic analysis using parsimony (* and other methods).. Sinauer Associates, Sunderland, Massachusetts.

Sytnikova, Y. A., R. Rahman, G.-w. Chirn, J. P. Clark, and N. C. Lau, 2014 Transposable element dynamics and PIWI regulation impacts lncRNA and gene expression diversity in Drosophila ovarian cell cultures. Genome Res 24: 1977–1990.

Vendrell-Mir, P., F. Barteri, M. Merenciano, J. González, J. M. Casacuberta, et al., 2019 A benchmark of transposon insertion detection tools using real data. Mob DNA 10: 53.

Wang, J., P. D. Keightley, and D. L. Halligan, 2007 Effect of divergence time and recombination rate on molecular evolution of Drosophila INE-1 transposable elements and other candidates for neutrally evolving sites. J Mol Evol 65: 627.

Wen, J., J. Mohammed, D. Bortolamiol-Becet, H. Tsai, N. Robine, et al., 2014 Diversity of miRNAs, siRNAs, and piRNAs across 25 Drosophila cell lines. Genome Res 24: 1236–1250.

Yanagawa, S.-i., J.-S. Lee, and A. Ishimoto, 1998 Identification and Characterization of a Novel Line ofDrosophila Schneider S2 Cells That Respond to Wingless Signaling. J Biol Chem 273: 32353–32359.

Yang, N., S. P. Srivastav, R. Rahman, Q. Ma, G. Dayama, et al., 2021 Transposable element landscape changes are buffered by RNA silencing in aging Drosophila. bioRxiv.

Yates, A. D., P. Achuthan, W. Akanni, J. Allen, J. Allen, et al., 2020 Ensembl 2020. Nucleic Acids Res 48: D682–D688.

Yu, M., S. K. Selvaraj, M. M. Y. Liang-Chu, S. Aghajani, M. Busse, et al., 2015 A resource for cell line authentication, annotation and quality control. Nature 520: 307–311.

Yu, T., X. Huang, S. Dou, X. Tang, S. Luo, et al., 2021 A benchmark and an algorithm for detecting germline transposon insertions and measuring de novo transposon insertion frequencies. Nucleic Acids Res.

Zaaijer, S., A. Gordon, D. Speyer, R. Piccone, S. C. Groen, et al., 2017 Rapid re-identification of human samples using portable DNA sequencing. eLife 6: e27798.

Zampella, J. G., N. Rodic, W. R. Yang, C. R. L. Huang, J. Welch, et al., 2016 A map of mobile DNA insertions in the NCI-60 human cancer cell panel. Mob DNA 7: 20.

Zanni, V., A. Eymery, M. Coiffet, M. Zytnicki, I. Luyten, et al., 2013 Distribution, evolution, and diversity of retrotransposons at the flamenco locus reflect the regulatory properties of piRNA clusters. Proc Natl Acad Sci USA 110.

Zhuang, J., J. Wang, W. Theurkauf, and Z. Weng, 2014 TEMP: a computational method for analyzing transposable element polymorphism in populations. Nucleic Acids Res 42: 6826– 6838.

