## Supplemental Text, Figures and Tables for "Transposable element profiles reveal cell line identity and loss of heterozygosity in *Drosophila* cell culture"

### 1 Supplementary Text

#### 1.1 Description of the ngs_te_mapper2 method for detecting non-reference TE insertions in single-end whole genome shotgun data

ngs_te_mapper2 (https://github.com/bergmanlab/ngs_te_mapper2) is a re-implementation of the method for detecting non-reference TE insertions in single-end whole genome shotgun sequence data initially reported in Linheiro and Bergman (2012). ngs_te_mapper2 uses a three-stage procedure to annotate non-reference TEs as the span of target site duplication (TSD) (Fig. S9), following the annotation framework described in Bergman (2012). In the first stage, whole genome shotgun (WGS) reads are mapped to a library of TE sequences to identify 'junction reads' that span the start/end of TE and genomic flanking sequences are retained. Such reads are often referred as 'split reads,' although in reality these reads are not split in the resequenced genome.

In the second stage, junction reads from each side of TE insertion identified in the first stage are separately mapped to a reference genome that has been hard-masked with RepeatMasker (http://www.repeatmasker.org/) using the same TE library from stage one (Fig. S9). Genome-wide coverage profiles are computed using samtools v1.9 (Li *et al.*, 2009) and genomic intervals with enriched coverage from junction read clusters on the 5' and 3' side of TEs are annotated in bed format. Regions of overlap between intervals of junction read clusters from the 5' and 3' side of TEs in the resequenced genome define the locations of TSDs for predicted non-reference TE insertions. The strand of non-reference TE predictions is determined from the relative orientation of alignments of the junction reads to the reference genome and TE library.

In the third stage, all reads from the original whole genome shotgun sequence data are mapped against the same hard-masked reference genome as in stage two (Fig. S9). This additional mapping step is necessary to obtain all reads that span the TE-flank junctions, as well as identify if any reads are present for the alternative "reference" haplotype that does not carry the non-reference TE insertion. For each candidate non-reference TE insertion site, the number of junction reads covering 5' and 3' side of each candidate TE insertion are estimated as the number of soft-clipped reads overlapping a 10bp window on the 5' and 3' side of the TSD, respectively ($\text{Count}_{junction5'}$ and $\text{Count}_{junction3'}$). The number of non-reference reads ($\text{Count}_{non-ref}$) is estimated as max($\text{Count}_{junction5'}$, $\text{Count}_{junction3'}$). The number of reference reads ($\text{Count}_{ref}$) is estimated as number of non-soft-clipped reads spanning the TSD with at least 3bp extension on both sides of the TSD. The allele frequency for non-reference TEs is heuristically estimated as $\text{Count}_{non-ref}/(\text{Count}_{non-ref} + \text{Count}_{ref})$.

#### 1.2 Evaluation of ngs_te_mapper2 performance

To evaluate the prediction performance of ngs_te_mapper2 and ngs_te_mapper under ideal conditions (one homozygous non-reference TE insertion with a known location), we created artificial ISO1 (dm6) genomes that each contain a single synthetic transposon insertion from one of the 125 TE families (excluding INE-1) in the Berkeley *Drosophila* Genome Project canonical TE dataset v10.1 (https://github.com/bergmanlab/transposons/blob/master/releases/D_mel_transposon_sequence_set_v10.1.fa; revision f94d53ea10b95c9da99258ac2336ce18871768e9). Insertion sites were selected at random in regions of normal recombination that were more than 500 bp from a reference TE in the *D. melanogaster* release 6.38 genome annotation (http://ftp.flybase.net/releases/FB2021_01/dmel_r6.38/gff/dmel-all-r6.38.gff.gz). After selecting an insertions site, a 5bp target site duplication was created and the full length canonical TE sequences was inserted into an otherwise unmodified dm6 genome sequence.

Ten synthetic genomes were created for each family in the *D. melanogaster* TE set, excluding the inactive INE-1 family, leading to total of 1250 synthetic genomes, each with a single non-reference TE insertion. 625 synthetic genomes contained a non-reference TE insertion of the TE canonical sequence (positive strand insertions), and 625 contained a non-reference TE insertion of the reverse complement of the TE canonical sequence (negative strand insertions). For each synthetic genome, 100 bp paired-end reads were simulated at 14X, 25X, 50X, and 100X coverage using wgsim v0.3.1-r13 ((Li, 2015), -e 0.01 -d 500). The forward reads of each simulated read pair, the unmodified dm6 reference genome, and the Berkeley *Drosophila* Genome Project canonical TE dataset v10.1 were used as input for ngs_te_mapper and ngs_te_mapper2 to detect non-reference TE insertions using McClintock (revision 40863acf11052b18afb4cdcd7b1124de48cba397; options: -m ngs_te_mapper, ngs_te_mapper2). Non-reference insertion predictions from ngs_te_mapper and ngs_te_mapper2 were considered a true positive if they occurred within 5bp of the actual synthetic insertion location and have the same TE family. Benchmark results under the single homozygous insertion scenario are summarized in Table S4. Under ideal conditions, the recall for ngs_te_mapper2 is high ($\geq$91.7%) and far exceeds that of ngs_te_mapper for all coverage levels. Likewise, in this idealized simulation setting the precision for ngs_te_mapper2 is $\geq$97.0% and the same as or better than ngs_te_mapper at all coverage levels.

Simulation of single homozygous insertion in unique regions of the dm6 reference genome provides a benchmark of ngs_te_mapper2 under ideal conditions, but does not incorporate the reality that TEs can insert into more complex regions of the genome, can exist in heterozygous state and are multiple TEs are predicted simultantously in real samples. To model both homozygous and heterozygous non-reference TE insertions and evaluate ngs_te_mapper2 under a more realistic setting, we created synthetic datasets using reads simulated from the ISO1 (dm6) and A4 (GCA_003401745.1) (Chakraborty *et al.*, 2018) genome assemblies. In theory, a good predictor should be able to accurately predict "non-reference" insertions that are present in genome 1 (e.g. ISO1) but absent from genome 2 (e.g. A4) using reads simulated from genome 1 mapped to genome 2. We therefore simulated 100bp synthetic paired-end sequencing data from the ISO1 genome assembly under 14X, 25X, 50X, 100X coverages using wgsim v0.3.1-r13 ((Li, 2015), -e 0.01 -d 500) to model homozygous insertions. Additionally, we simulated synthetic paired-end sequencing data by combining equal numbers of reads from both ISO1 and A4 genome assemblies to model heterozygous insertions. The synthetic datasets were used as input to ngs_te_mapper2 to detect non-reference TE insertions using McClintock (revision 40863acf11052b18afb4cdcd7b1124de48cba397; options: -m "trimgalore, ngs_te_mapper2, map_reads"). The A4 assembly was used as the reference genome and the Berkeley *Drosophila* Genome Project canonical TE dataset v10.1 were used for these analyses.

As ground truth for evaluating ngs_te_mapper2 performance, curated TE annotations from the release 6.38 version of *D. melanogaster* genome (http://ftp.flybase.net/releases/FB2021_01/dmel_r6.38/gff/dmel-all-r6.38.gff.gz) were lifted over to A4 genome assembly. After excluding INE-1 insertions and TE insertions in low recombination regions, 627 curated TEs in ISO1 could be lifted over to A4 on the basis of their flanking regions. ngs_te_mapper2 predictions were considered true positives if the predicted TE insertion coordinates were within a 5bp window of a lifted over ISO1 TE annotation and if the predicted TE family was the same as the lifted over annotation. The final benchmark results for ngs_te_mapper2 applied to simulated real genomes are summarized in Table S5. Similar to single synthetic insertion simulations above, ngs_te_mapper2 has high precision ($\geq$95.0%) at all coverage levels in simulations designed to model genome-wide TE prediction. In contrast, recall for ngs_te_mapper2 under a more realistic setting was much lower than in single synthetic insertion simulations, especially at low coverage levels, and was lower for heterozygous insertions than homozygous insertions at all coverage levels. These results indicate that the TE insertion predictions ngs_te_mapper2 makes are accurate but that the method has an appreciable false negative rate on low coverage samples.

#### 1.3 Evaluation of a classifier for predicting homozygous or heterozygous TE insertion in single-end WGS data

To fill a gap in tools available to analyze intra-sample TE allele frequencies in single-end WGS data, we developed a classifier to determine whether a TE insertion predicted by ngs_te_mapper2 is homozygous or heterozygous. Our model classifies a TE insertion as homozygous if the intra-sample allele frequency is $\geq$0.95, as heterozygous if the allele frequency is between 0.25 and 0.75, and is considered unclassified if neither of these conditions are met. To evaluate this approach we used ngs_te_mapper2 predictions made from the simulated paired-end sequencing data generated from ISO1 and A4 genome assemblies described in the previous section. We evaluated the classifier as follows: if the simulated reads were generated from ISO1 only, then all all non-reference TE insertions were expected to be homozygous and the precision was calculated as $\text{Count}_{homozygous}$ / $\text{Count}_{all}$. If the simulated data were a combination of reads from both ISO1 and A4, then all non-reference TE insertions were expected to be heterozygous and the precision is $\text{Count}_{heterozygous}$ / $\text{Count}_{all}$. The final benchmark results were summarized in Table S6. Our classifier had $\geq$91.3% precision at all coverage levels and never falsely classified a heterozygous TE insertions as homozygous, and is thus conservative for the purposes of detecting loss of heterozygosity.

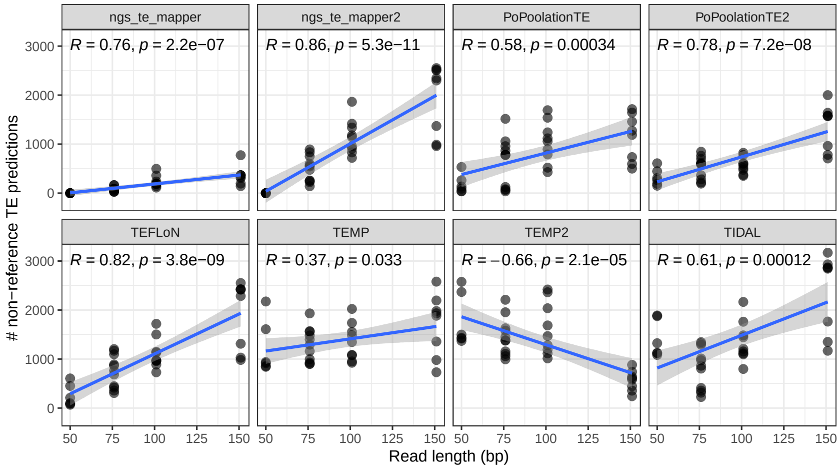


**Figure S1. Relationship between read length and number of non-reference TE predictions for the expanded dataset of 34 *Drosophila* cell line samples.** Each panel represents predictions from one of the eight component methods designed for detection of TE insertions in *Drosophila* that is included in McClintock. The X-axis represents read length in base pairs (bp) and the Y-axis represents the number of non-reference TE predictions. The best fit line and 95% CI were included using linear method. Pearson correlation coefficient with p-values are shown on the top of each panel.

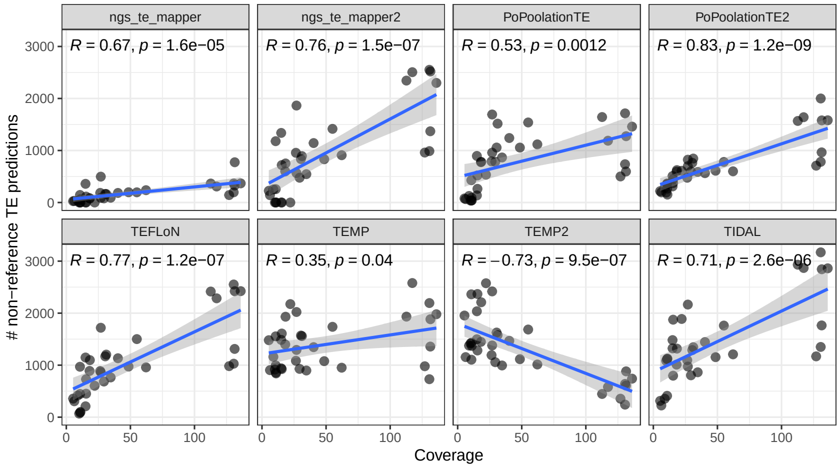


**Figure S2. Relationship between average genome coverage and number of non-reference TE predictions for the expanded dataset of 34 *Drosophila* cell line samples.** Each panel represents predictions from one of the eight component methods designed for detection of TE insertions in *Drosophila* that is included in McClintock. The X-axis represents the average genome coverage computed by McClintock and the Y-axis represents the number of non-reference TE predictions. The best fit line and 95% CI were included using linear method. Pearson correlation coefficient with p-values are shown on the top of each panel.

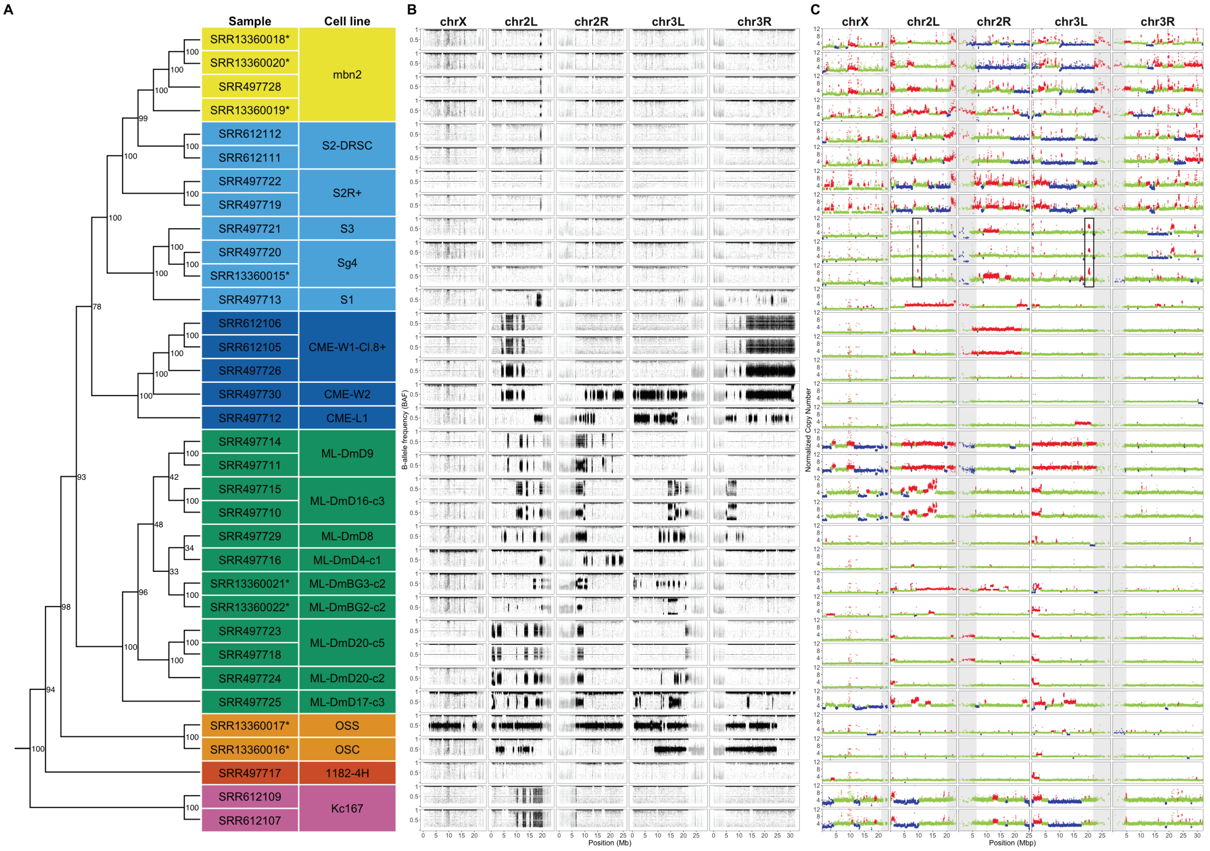


**Figure S3. Copy number and B-allele frequency profiles for the expanded dataset of 34 *Drosophila* cell line samples** (A) Dollo parsimony tree of 34 *Drosophila* cell lines samples (including replicates and sub-lines) based on non-reference TE predictions. Node labels indicate support for each clade based on 100 bootstrap replicates. New sequence data from this study are indicated by asterisks. (B) B-allele frequency profiles for *Drosophila* cell lines on major chromosome arms. For a given SNP, the B-allele frequency (BAF) was determined as the coverage of reads supporting non-reference allele divided by total coverage at that position. SNPs in low recombination regions are plotted in grey. (C) Copy number profiles for *Drosophila* cell lines on major chromosome arms. Each data point represents normalized copy number (ratio*ploidy) for a given 10kb window estimated by Control-FREEC (Boeva *et al.*, 2012). Data points for each window are colorized by CNV status (red: CNV gain; green: no CNV; blue: CNV loss), which are based on the comparison between normalized copy number for that window and baseline ploidy for the chromosome arm. Black boxes in panel C highlight regions where Sg4 and S3 cell lines share the same copy number gains that are not shared in other cell samples. Low recombination regions are shaded in grey.

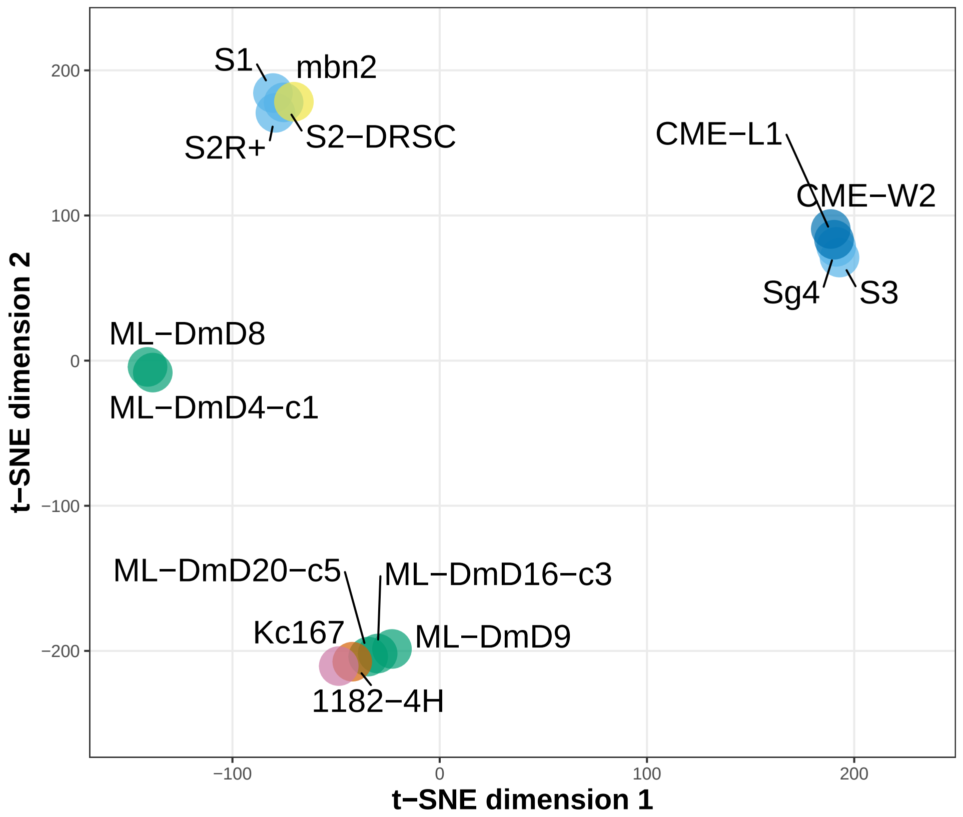


**Figure S4. t-SNE visualization of 15 *Drosophila* cell lines using total RNA-seq data from Brown *et al.* (2014).** t-SNE visualization was produced with perplexity=1. Samples are colorized by the lab origin of cell lines.

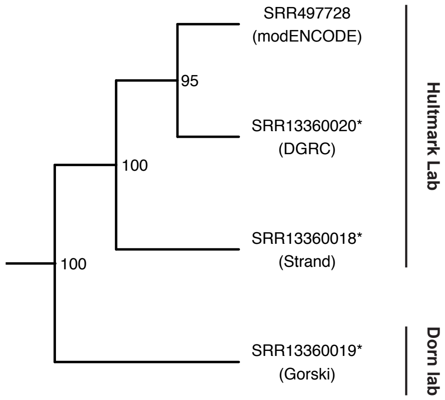


**Figure S5. Clustering of normalized mbn2 cell line genome samples from the modENCODE project plus this study.** Clustering was performed on TE insertions generated using mbn2 samples that were normalized by trimming read lengths to 76bp and downsampling to 19x depth. For this analysis, we also relaxed TEMP filtering to include more weakly-supported predictions at otherwise high-quality loci because of the lower overall coverage in all samples. Numbers beside nodes indicate percent support based on 100 bootstrap replicates. Tip labels include SRA run identifiers and source lab for samples (in parentheses). New sequence data from this study are indicated by asterisks. Clade annotations indicate the donor lab from which the source lab obtained their sub-line of mbn2 cells.

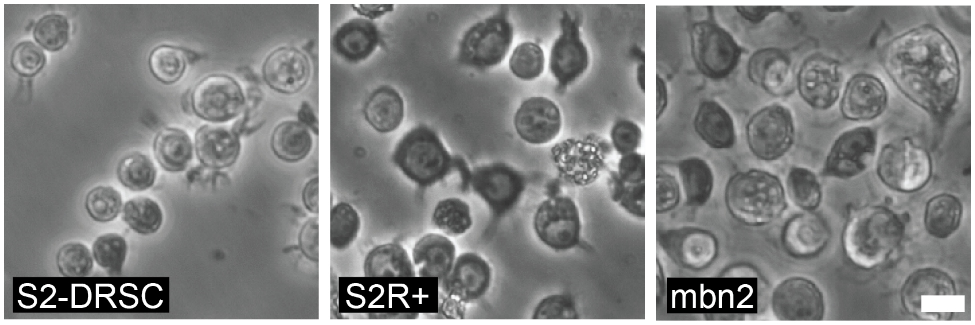


**Figure S6. Morphology of S2, S2R+ and mbn2 cell lines** Phase-contrast micrographs of S2-DRSC (DGRC-181), S2R+ (DGRC-150), and mbn2 (DGRC-147). Scale bar is 10 microns.

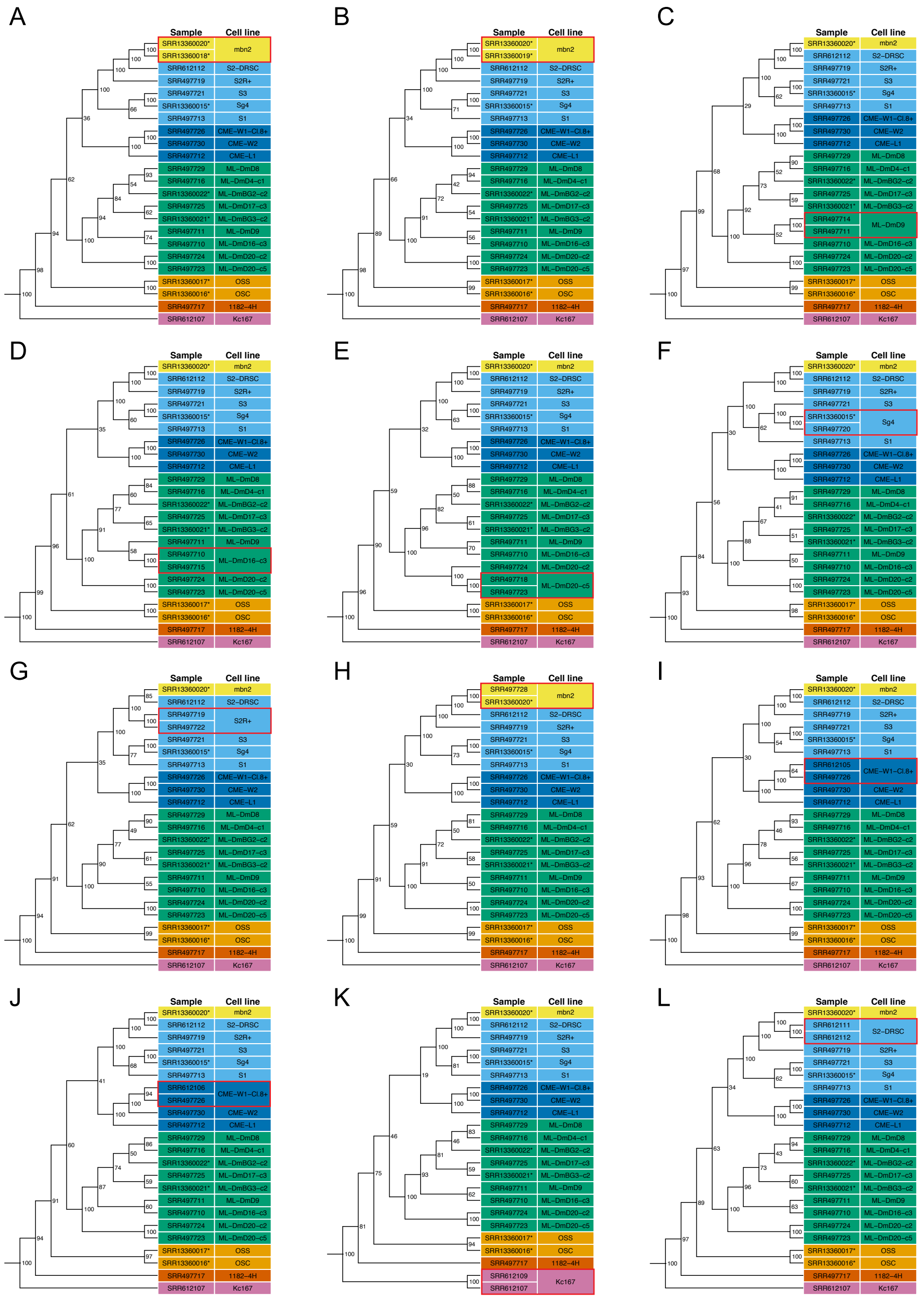


**Figure S7. *Drosophila* cell line samples can be identified using TE profiles from a diagnostic set of six LTR retrotransposon families.** Panels represent Dollo parsimony trees of a common set of 22 *Drosophila* cell line primary replicates plus one additional secondary replicate, one tree for each of the 12 secondary replicates from the nine cell lines in the expanded dataset with secondary replicates. Dollo parsimony trees were constructed using non-reference TE predictions for six *D. melanogaster* LTR retrotransposon families (297, copia, mdg3, mdg1, roo and 1731). Samples are colorized by lab origin. Cell lines with secondary replicates are highlighted in red boxes. Node labels indicate support for each clade based on 100 bootstrap replicates. New sequence data from this study are indicated by asterisks.

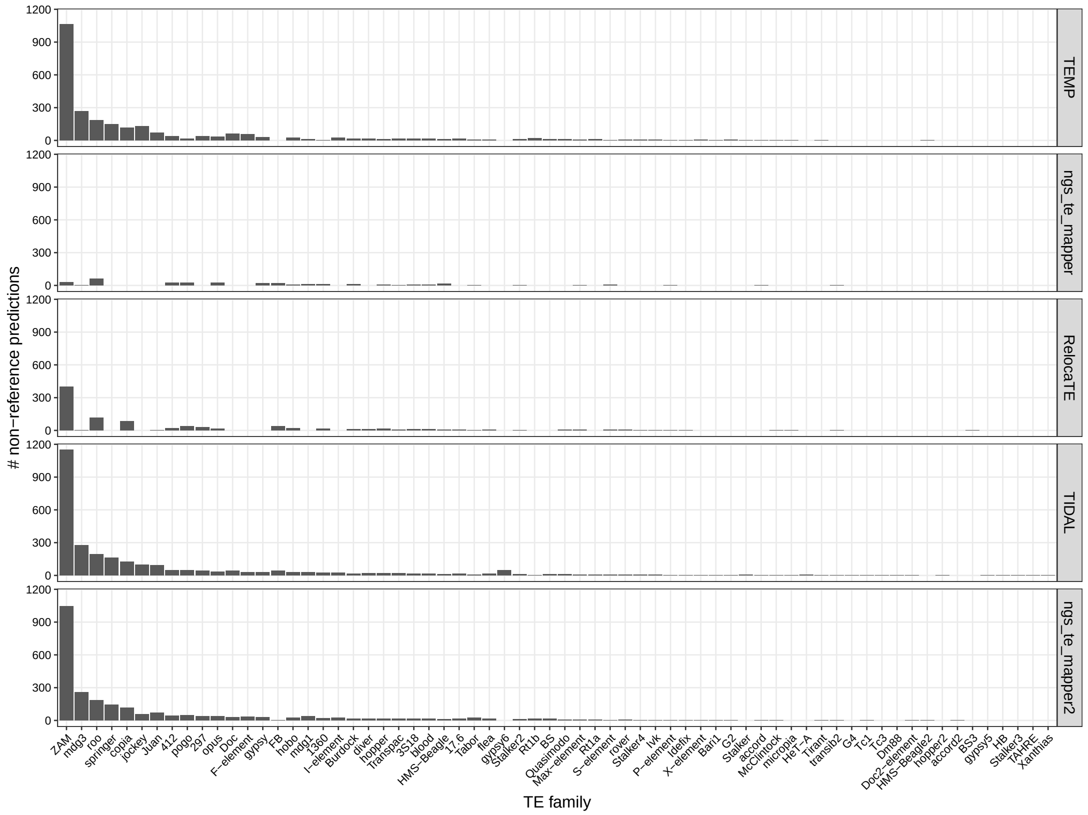


**Figure S8. Number of non-reference TE predictions on trimmed downsampled OSS_DGRC using five TE detection methods.** Paired-end sequencing data for OSS_DGRC was used as input for TEMP, ngs_te_mapper, RelocaTE, TIDAL and ngs_te_mapper2 to detect non-reference TE insertions using McClintock. INE-1 and insertion predictions in low recombination regions were excluded from all panels.

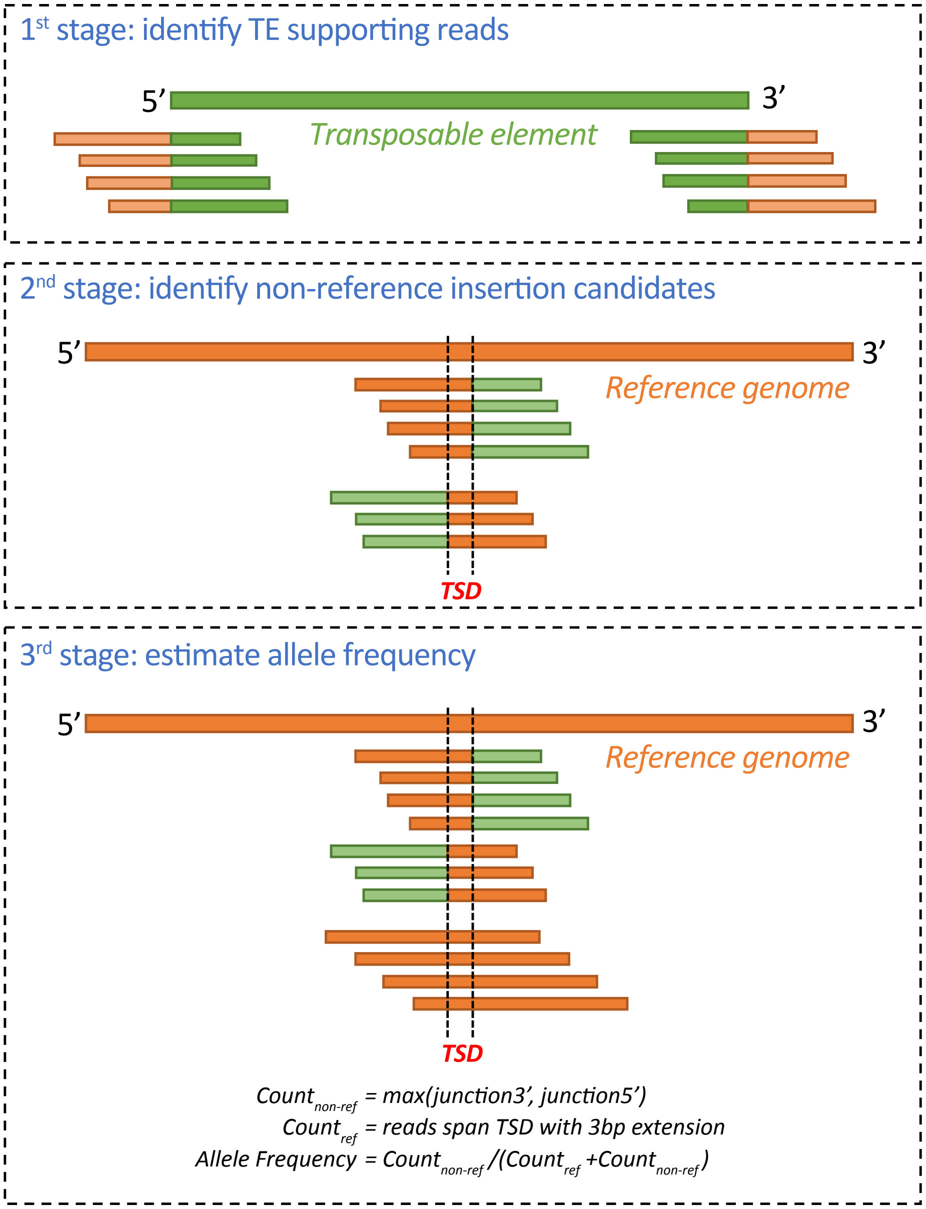


**Figure S9. ngs_te_mapper2 workflow for predicting non-reference TE insertions.** In the first stage, raw reads are mapped to the TE consensus sequences. Reads that partially map to TEs are extracted as putative TE supporting reads. In the second stage, putative TE supporting reads are mapped to reference genome that has been hard-masked with RepeatMasker using input TE library. Non-reference TE insertion candidates are identified if alignments of TE supporting reads on 5' and 3' end of TE overlap. In the third stage, raw reads are mapped to unmodified reference genome for estimating intra-sample insertion allele frequency. See details in section 1.1.

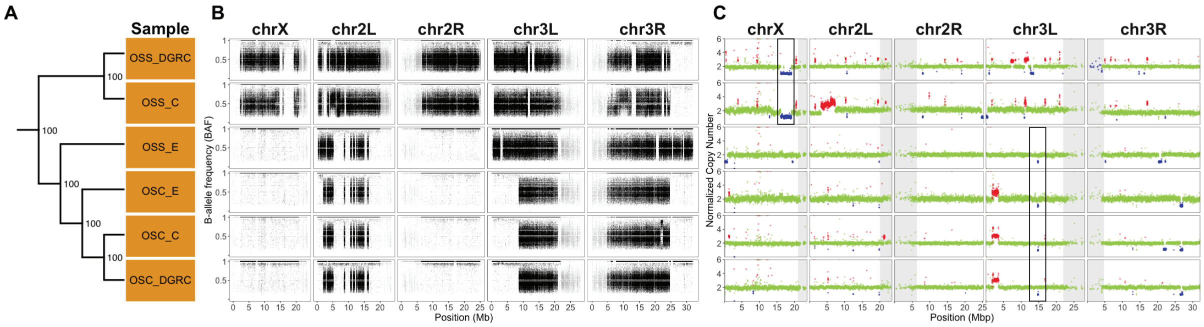


**Figure S10. Copy number and B-allele frequency profiles for six ovarian cell line samples** (A) Dollo parsimony tree of six ovarian cell line samples based on non-reference TE predictions excluding ZAM insertions using single-end WGS data. Node labels indicate support for each clade based on 100 bootstrap replicates. (B) B-allele frequency profiles for ovarian cell line samples on major chromosome arms. For a given SNP, the B-allele frequency (BAF) was determined as the coverage of reads supporting non-reference allele divided by total coverage at that position. SNPs in low recombination regions are plotted in grey. (C) Copy number profiles for ovarian cell line samples on major chromosome arms. Each data point represents normalized copy number (ratio*ploidy) for a given 10kb window estimated by Control-FREEC (Boeva *et al.*, 2012). Data points for each window are colorized by CNV status (red: CNV gain; green: no CNV; blue: CNV loss), which are based on the comparison between normalized copy number for that window and baseline ploidy for the chromosome arm. Black boxes in panel C highlight regions where cell lines share the same copy number loss events that are not shared in other cell samples. Low recombination regions are shaded in grey.

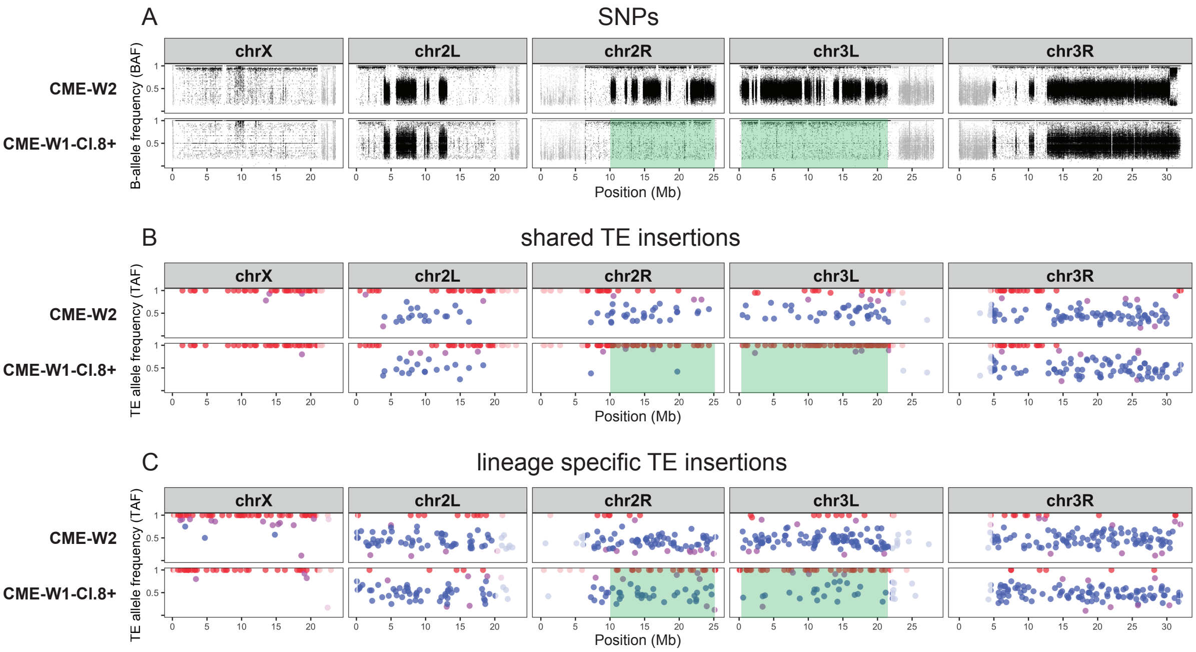


**Figure S11. Loss of heterozygosity and ongoing transposition shape TE profiles in *Drosophila* imaginal disc derived cell lines.** Allele frequency profiles for CME-W2 and CME-W1-Cl.8+ cell lines based on (A) SNP variants, (B) TE insertions shared by CME-W2 and CME-W1-Cl.8+, and (C) lineage specific TE insertions restricted to only CME-W2 or CME-W1-Cl.8+. SNPs and TE insertions in highly-repetitive low recombination regions are shaded in grey. TE insertions are classified as being homozygous (red), heterozygous (blue), or undefined (purple) based on allele frequencies estimated by ngs_te_mapper2. Green shading indicates LOH regions defined by the more extensive pattern of SNP heterozygosity in CME-W2 relative to CME-W1-Cl.8+.

**Table S1. Metadata and sequencing information for 34 paired-end whole genome shotgun sequencing samples from 22 *Drosophila* cell lines used in this study.** Samples indicated by an asterisk were generated in the current study, while other samples were generated by the modENCODE project (Lee *et al.*, 2014). *Drosophila* Genomics Resource Center (DGRC) cell line names and stock identifiers are given for all cell line samples except two mbn2 samples obtained from the Gorski lab (Canada's Michael Smith Genome Sciences Centre, BC Cancer) and the Strand lab (University of Georgia), respectively. For DGRC cell lines, the donor lab represents the lab who donated the stock to the DGRC. Ancestral genotypes represents the genotype of flies from which the cell lines were established. Inferred ploidy represents the ploidy estimated by analyzing DNA density of whole genome data using the method of Lee *et al.* (2014). Inferred sex represents the sex of the cell line inferred by analyzing DNA density of whole genome data and analysis of sex determination gene expression based on Lee *et al.* (2014). Coverage represents the average mapped depth of coverage after quality and adaptor trimming. N.A. indicates that this information is not available.

| Cell line | DGRC ID | FlyBase ID | Donor lab | Lab origin | Ancestral genotype | Inferred ploidy | Inferred sex | SRA | Read length | Coverage | Primary replicate |
|---|---|---|---|---|---|---|---|---|---|---|---|
| 1182-4H | DGRC-177 | FBtc0000177 | Debec | Debec | mh | 2 | female | SRR497717 | 101 | 26.46 | yes |
| CME-L1 | DGRC-156 | FBtc0000156 | Cottam & Milner | Milner | Oregon-R | 2 | male | SRR497712 | 101 | 62.17 | yes |
| CME-W1-Cl.8+ | DGRC-151 | FBtc0000151 | Cottam & Milner | Milner | Oregon-R | 2 | male | SRR612105 | 50 | 10.99 | no |
| CME-W1-Cl.8+ | DGRC-151 | FBtc0000151 | Cottam & Milner | Milner | Oregon-R | 2 | male | SRR612106 | 50 | 10.05 | no |
| CME-W1-Cl.8+ | DGRC-151 | FBtc0000151 | Cottam & Milner | Milner | Oregon-R | 2 | male | SRR497726 | 76 | 18.14 | yes |
| CME-W2 | DGRC-155 | FBtc0000155 | Cottam & Milner | Milner | Oregon-R | 2 | male | SRR497730 | 76 | 31.15 | yes |
| Kc167 | DGRC-1 | FBtc0000001 | Cherbas | Echalier | e/se | 4 | female | SRR612107 | 50 | 15.01 | yes |
| Kc167 | DGRC-1 | FBtc0000001 | Cherbas | Echalier | e/se | 4 | female | SRR612109 | 50 | 10.82 | no |
| mbn2 | DGRC-147 | FBtc0000147 | Werner & Hultmark | Gateff | l(2)mbn | 4 | male | SRR497728 | 76 | 18.38 | no |
| mbn2 (*) | DGRC-147 | FBtc0000147 | Werner & Hultmark | Gateff | l(2)mbn | 4 | male | SRR13360020 | 151 | 112.69 | yes |
| mbn2 (Gorski) (*) | N.A. | N.A. | Gorski | Gateff | l(2)mbn | 4 | male | SRR13360019 | 151 | 130.60 | no |
| mbn2 (Strand) (*) | N.A. | N.A. | Strand | Gateff | l(2)mbn | 4 | male | SRR13360018 | 151 | 136.06 | no |
| ML-DmBG2-c2 (*) | DGRC-53 | FBtc0000053 | Ueda & Ui-Tei | Miyake | $y^1 v^1 f^1 mal^{F1}$ | 2 | male | SRR13360022 | 151 | 127.03 | yes |
| ML-DmBG3-c2 (*) | DGRC-68 | FBtc0000068 | Ueda & Ui-Tei | Miyake | $y^1 v^1 f^1 mal^{F1}$ | 2 | male | SRR13360021 | 151 | 130.59 | yes |
| ML-DmD16-c3 | DGRC-97 | FBtc0000097 | Ueda & Ui-Tei | Miyake | $y^1 v^1 f^1 mal^{F1}$ | 4 | female | SRR497715 | 76 | 10.73 | no |
| ML-DmD16-c3 | DGRC-97 | FBtc0000097 | Ueda & Ui-Tei | Miyake | $y^1 v^1 f^1 mal^{F1}$ | 4 | female | SRR497710 | 101 | 48.55 | yes |
| ML-DmD17-c3 | DGRC-107 | FBtc0000107 | Ueda & Ui-Tei | Miyake | $y^1 v^1 f^1 mal^{F1}$ | 4 | female | SRR497725 | 101 | 55.02 | yes |
| ML-DmD20-c2 | DGRC-109 | FBtc0000109 | Ueda & Ui-Tei | Miyake | $y^1 v^1 f^1 mal^{F1}$ | 2 | male | SRR497724 | 76 | 26.93 | yes |
| ML-DmD20-c5 | DGRC-112 | FBtc0000112 | Ueda & Ui-Tei | Miyake | $y^1 v^1 f^1 mal^{F1}$ | 2 | male | SRR497718 | 76 | 6.24 | no |
| ML-DmD20-c5 | DGRC-112 | FBtc0000112 | Ueda & Ui-Tei | Miyake | $y^1 v^1 f^1 mal^{F1}$ | 2 | male | SRR497723 | 101 | 15.42 | yes |
| ML-DmD4-c1 | DGRC-126 | FBtc0000126 | Ueda & Ui-Tei | Miyake | $y^1 v^1 f^1 mal^{F1}$ | 2 | male | SRR497716 | 76 | 34.62 | yes |
| ML-DmD8 | DGRC-92 | FBtc0000092 | Ueda & Ui-Tei | Miyake | $y^1 v^1 f^1 mal^{F1}$ | 2 | female | SRR497729 | 76 | 29.34 | yes |
| ML-DmD9 | DGRC-85 | FBtc0000085 | Ueda & Ui-Tei | Miyake | $y^1 v^1 f^1 mal^{F1}$ | 4 | female | SRR497714 | 76 | 8.89 | no |
| ML-DmD9 | DGRC-85 | FBtc0000085 | Ueda & Ui-Tei | Miyake | $y^1 v^1 f^1 mal^{F1}$ | 4 | female | SRR497711 | 101 | 40.28 | yes |
| OSC (*) | DGRC-288 | FBtc0000288 | Saito & Siomi | Niki | w1118 | 2 | female | SRR13360016 | 151 | 131.31 | yes |
| OSS (*) | DGRC-190 | FBtc0000190 | Niki | Niki | w1118 | 2 | female | SRR13360017 | 151 | 117.27 | yes |
| S1 | DGRC-9 | FBtc0000009 | Cherbas | Schneider | Oregon-R | 2 | male | SRR497713 | 76 | 30.39 | yes |
| S2-DRSC | DGRC-181 | FBtc0000181 | Perrimon & Mathey-Prevot | Schneider | Oregon-R | 4 | male | SRR612111 | 50 | 15.45 | no |
| S2-DRSC | DGRC-181 | FBtc0000181 | Perrimon & Mathey-Prevot | Schneider | Oregon-R | 4 | male | SRR612112 | 50 | 22.10 | yes |
| S2R+ | DGRC-150 | FBtc0000150 | Wheeler | Schneider | Oregon-R | 4 | male | SRR497722 | 76 | 5.32 | no |
| S2R+ | DGRC-150 | FBtc0000150 | Wheeler | Schneider | Oregon-R | 4 | male | SRR497719 | 101 | 10.66 | yes |
| S3 | DGRC-5 | FBtc0000005 | Cherbas | Schneider | Oregon-R | 4 | male | SRR497721 | 101 | 14.99 | yes |
| Sg4 | DGRC-179 | FBtc0000179 | Pirrotta | Schneider | Oregon-R | 4 | male | SRR497720 | 101 | 26.91 | no |
| Sg4 (*) | DGRC-179 | FBtc0000179 | Pirrotta | Schneider | Oregon-R | 4 | male | SRR13360015 | 151 | 131.41 | yes |

**Table S2. Summary of predictions generated by eight non-reference TE insertion detection methods for 34 *Drosophila* cell line samples.** Numbers of non-reference TE insertion predictions are based on default settings for TIDAL (Rahman *et al.*, 2015; Yang *et al.*, 2021) and default McClintock (Nelson *et al.*, 2017) settings for all other methods. INE-1 and non-reference TE insertion predictions in low recombination regions were excluded from all methods. New sequence data from this study are indicated by asterisks.

| Cell line | SRA | TEMP | TEMP2 | PoPoolationTE | PoPoolationTE2 | TEFLoN | ngs_te_mapper | ngs_te_mapper2 | TIDAL |
|---|---|---|---|---|---|---|---|---|---|
| 1182-4H | SRR497717 | 1084 | 1192 | 790 | 476 | 887 | 185 | 956 | 1096 |
| CME-L1 | SRR497712 | 951 | 1013 | 1118 | 600 | 957 | 237 | 909 | 1208 |
| CME-W1-Cl.8+ | SRR612105 | 841 | 1370 | 40 | 152 | 100 | 0 | 1 | 1125 |
| CME-W1-Cl.8+ | SRR612106 | 915 | 1418 | 37 | 192 | 68 | 0 | 0 | 1083 |
| CME-W1-Cl.8+ | SRR497726 | 1398 | 1448 | 776 | 594 | 889 | 83 | 599 | 1010 |
| CME-W2 | SRR497730 | 1559 | 1578 | 1516 | 845 | 1203 | 163 | 892 | 1345 |
| Kc167 | SRR612107 | 939 | 1498 | 136 | 309 | 209 | 0 | 0 | 1323 |
| Kc167 | SRR612109 | 856 | 1432 | 91 | 266 | 88 | 0 | 0 | 1119 |
| mbn2 | SRR497728 | 1931 | 2209 | 781 | 623 | 1097 | 77 | 751 | 1318 |
| mbn2 (*) | SRR13360020 | 1933 | 446 | 1643 | 1568 | 2415 | 365 | 2344 | 2931 |
| mbn2 (Gorski) (*) | SRR13360019 | 2194 | 639 | 1714 | 2000 | 2551 | 366 | 2551 | 3169 |
| mbn2 (Strand) (*) | SRR13360018 | 1979 | 740 | 1459 | 1581 | 2423 | 368 | 2299 | 2862 |
| ML-DmBG2-c2 (*) | SRR13360022 | 979 | 355 | 501 | 708 | 981 | 143 | 960 | 1169 |
| ML-DmBG3-c2 (*) | SRR13360021 | 730 | 241 | 736 | 782 | 1028 | 196 | 988 | 1350 |
| ML-DmD16-c3 | SRR497715 | 995 | 1105 | 37 | 365 | 446 | 57 | 257 | 410 |
| ML-DmD16-c3 | SRR497710 | 1077 | 1115 | 1056 | 611 | 973 | 197 | 832 | 1154 |
| ML-DmD17-c3 | SRR497725 | 1737 | 1685 | 1539 | 780 | 1501 | 196 | 1416 | 1761 |
| ML-DmD20-c2 | SRR497724 | 1293 | 1379 | 958 | 699 | 864 | 89 | 566 | 973 |
| ML-DmD20-c5 | SRR497718 | 904 | 1155 | 65 | 195 | 306 | 30 | 141 | 225 |
| ML-DmD20-c5 | SRR497723 | 924 | 1282 | 520 | 374 | 729 | 114 | 720 | 797 |
| ML-DmD4-c1 | SRR497716 | 897 | 994 | 866 | 588 | 764 | 92 | 545 | 863 |
| ML-DmD8 | SRR497729 | 928 | 1057 | 783 | 596 | 685 | 80 | 476 | 805 |
| ML-DmD9 | SRR497714 | 1160 | 1376 | 126 | 283 | 416 | 34 | 252 | 354 |
| ML-DmD9 | SRR497711 | 1346 | 1468 | 1240 | 564 | 1133 | 184 | 1143 | 1444 |
| OSC (*) | SRR13360016 | 1357 | 607 | 596 | 964 | 1312 | 327 | 1370 | 1764 |
| OSS (*) | SRR13360017 | 2579 | 580 | 1188 | 1640 | 2285 | 308 | 2506 | 2868 |
| S1 | SRR497713 | 1569 | 1627 | 1057 | 763 | 1169 | 165 | 835 | 1290 |
| S2-DRSC | SRR612111 | 1608 | 2368 | 263 | 447 | 451 | 0 | 0 | 1874 |
| S2-DRSC | SRR612112 | 2174 | 2575 | 534 | 609 | 604 | 0 | 0 | 1888 |
| S2R+ | SRR497722 | 1480 | 1954 | 78 | 219 | 357 | 26 | 225 | 313 |
| S2R+ | SRR497719 | 1554 | 2362 | 429 | 353 | 969 | 145 | 1179 | 1114 |
| S3 | SRR497721 | 1486 | 2036 | 895 | 509 | 1147 | 359 | 1338 | 1482 |
| Sg4 | SRR497720 | 2022 | 2418 | 1692 | 823 | 1719 | 497 | 1864 | 2165 |
| Sg4 (*) | SRR13360015 | 1883 | 881 | 1275 | 1579 | 2421 | 774 | 2519 | 2844 |

**Table S3. Summary of transcriptome data for *Drosophila* cell lines analyzed in this study.** Samples are from two consistent batches of RNA-seq experiments performed on DGRC cell lines with genome data. The first batch is poly-A RNA-seq samples from Stoiber *et al.* (2016) (PRJNA306537) and the other batch is total RNA-seq samples from Brown *et al.* (2014) (PRJNA75285). All samples have 100 bp paired end reads.

| Cell_line | SRA | Study_accession | Gigabases |
|---|---|---|---|
| 1182-4H | SRR1197409 | PRJNA75285 | 10.0 |
| 1182-4H | SRR3038250 | PRJNA306537 | 3.7 |
| CME-L1 | SRR1197410 | PRJNA75285 | 10.8 |
| CME-L1 | SRR3038125 | PRJNA306537 | 4.4 |
| CME-W1-Cl.8+ | SRR3038123 | PRJNA306537 | 3.1 |
| CME-W2 | SRR1197407 | PRJNA75285 | 10.9 |
| CME-W2 | SRR3038127 | PRJNA306537 | 2.6 |
| Kc167 | SRR1197456 | PRJNA75285 | 11.6 |
| Kc167 | SRR3040509 | PRJNA306537 | 3.4 |
| mbn2 | SRR1197406 | PRJNA75285 | 9.3 |
| mbn2 | SRR3040560 | PRJNA306537 | 2.7 |
| ML-DmD16-c3 | SRR1197401 | PRJNA75285 | 10.1 |
| ML-DmD17-c3 | SRR3041988 | PRJNA306537 | 1.8 |
| ML-DmD20-c5 | SRR1197396 | PRJNA75285 | 10.4 |
| ML-DmD20-c5 | SRR3042157 | PRJNA306537 | 2.8 |
| ML-DmD4-c1 | SRR1197397 | PRJNA75285 | 10.3 |
| ML-DmD4-c1 | SRR3042204 | PRJNA306537 | 2.7 |
| ML-DmD8 | SRR1197284 | PRJNA75285 | 8.0 |
| ML-DmD8 | SRR3042539 | PRJNA306537 | 4.3 |
| ML-DmD9 | SRR1197283 | PRJNA75285 | 10.3 |
| ML-DmD9 | SRR3042543 | PRJNA306537 | 3.4 |
| S1 | SRR1197281 | PRJNA75285 | 8.8 |
| S1 | SRR3042563 | PRJNA306537 | 3.4 |
| S2-DRSC | SRR1197282 | PRJNA75285 | 9.6 |
| S2-DRSC | SRR3042565 | PRJNA306537 | 3.1 |
| S2R+ | SRR1197280 | PRJNA75285 | 9.0 |
| S3 | SRR1197277 | PRJNA75285 | 8.9 |
| S3 | SRR3042571 | PRJNA306537 | 3.8 |
| Sg4 | SRR1197278 | PRJNA75285 | 8.8 |
| Sg4 | SRR3042573 | PRJNA306537 | 4.9 |

**Table S4. ngs_te_mapper2 performance benchmark using single insertion synthetic data.** ngs_te_mapper (Linheiro and Bergman, 2012) and ngs_te_mapper2 were benchmarked by creating single synthetic TE insertions in the ISO1 (dm6) genome assembly, simulating reads from these modified assemblies under different coverages, then generating insertion predictions using unmodified assembly as reference genome and comparing predictions with expected insertion annotations. 10 single synthetic insertion simulation experiments were performed for each of the 125 TE families in *D. melanogaster* (excluding INE-1), making up 1250 total simulations each with one synthetic insertion. "Total" represents the total number of predictions from all 1250 experiments after filtering (see filtering criteria in section 1.2). "True Positives" and "False Positives" represent the number of predictions that match or don't match expected insertion annotations, respectively (see matching criteria in section 1.2). "False Negatives" represent the number of expected insertion annotations that are not predicted by the TE detection method. "Precision" represents the number of true positives divided by total number of predictions. "Recall" represents the number of true positives divided by total number of expected insertions (1250).

| Method | Coverage | Total | True Positives | False Positives | False Negatives | Precision | Recall |
|---|---|---|---|---|---|---|---|
| ngs_te_mapper | 14 | 487 | 481 | 6 | 769 | 98.8% | 38.5% |
| ngs_te_mapper | 25 | 525 | 519 | 6 | 731 | 98.9% | 41.5% |
| ngs_te_mapper | 50 | 534 | 527 | 7 | 723 | 98.7% | 42.2% |
| ngs_te_mapper | 100 | 533 | 524 | 9 | 726 | 98.3% | 41.9% |
| ngs_te_mapper2 | 14 | 1149 | 1146 | 3 | 104 | 99.7% | 91.7% |
| ngs_te_mapper2 | 25 | 1181 | 1173 | 8 | 77 | 99.3% | 93.8% |
| ngs_te_mapper2 | 50 | 1189 | 1174 | 15 | 76 | 98.7% | 93.9% |
| ngs_te_mapper2 | 100 | 1211 | 1175 | 36 | 75 | 97.0% | 94.0% |

**Table S5. ngs_te_mapper2 performance benchmark using genome-wide synthetic data from ISO1 and A4 genome assemblies.** Non-reference TE insertion predictions made by ngs_te_mapper2 using the A4 genome assembly as reference were evaluated against curated TE annotations in ISO1 lifted over to A4 coordinates (see section 1.2 for details). Zygosity represents whether simulated reads were generated from both ISO1 and A4 (heterozygous) or ISO1 only (homozygous). "True Positives" and "False Positives" represent the number of predictions that match and doesn't match with lifted over insertion annotations, respectively. "False Negatives" represent the number of lifted over non-reference TE insertion annotations that are not predicted by ngs_te_mapper2. "Precision" represents the number of true positives divided by total number of predictions. "Recall" represents the number of true positives divided by total number of lifted over non-reference TE insertion annotations.

| Zygosity | Coverage | Total | True positives | False positives | False negatives | Precision | Recall |
|---|---|---|---|---|---|---|---|
| heterozygous | 14 | 346 | 336 | 10 | 285 | 97.1% | 53.8% |
| heterozygous | 25 | 424 | 412 | 12 | 209 | 97.2% | 66.0% |
| heterozygous | 50 | 476 | 462 | 14 | 159 | 97.1% | 74.0% |
| heterozygous | 100 | 482 | 464 | 18 | 157 | 96.3% | 74.4% |
| homozygous | 14 | 437 | 424 | 13 | 197 | 97.0% | 67.9% |
| homozygous | 25 | 473 | 461 | 12 | 160 | 97.5% | 73.9% |
| homozygous | 50 | 482 | 465 | 17 | 156 | 96.5% | 74.5% |
| homozygous | 100 | 516 | 490 | 26 | 131 | 95.0% | 78.5% |

**Table S6. Performance benchmark for intra-sample TE insertion zygosity classifier.** ngs_te_mapper2 predictions on synthetic data from ISO1 and A4 genome assemblies were used as input for the classifier. Zygosity represents whether the simulated reads were generated from both ISO1 and A4 (heterozygous) or ISO1 only (homozygous). Precision represents the proportion of predictions being correctly classified as heterozygous or homozygous by the classifier.

| Zygosity | Coverage | Total | Homozygous count | Heterozygous count | Unclassified count | Precision |
|---|---|---|---|---|---|---|
| heterozygous | 14 | 346 | 0 | 326 | 20 | 94.2% |
| heterozygous | 25 | 424 | 0 | 419 | 5 | 98.8% |
| heterozygous | 50 | 476 | 0 | 473 | 3 | 99.4% |
| heterozygous | 100 | 482 | 0 | 477 | 5 | 99.0% |
| homozygous | 14 | 437 | 399 | 4 | 34 | 91.3% |
| homozygous | 25 | 473 | 438 | 3 | 32 | 92.6% |
| homozygous | 50 | 482 | 456 | 3 | 23 | 94.6% |
| homozygous | 100 | 516 | 489 | 9 | 18 | 94.8% |

### Supplemental References


Bergman, C. M., 2012 A proposal for the reference-based annotation of de novo transposable element insertions. Mob Genet Elements **2**: 51–54.

Boeva, V., T. Popova, K. Bleakley, P. Chiche, J. Cappo, *et al.*, 2012 Control-FREEC: a tool for assessing copy number and allelic content using next-generation sequencing data. Bioinformatics **28**: 423–425.

Brown, J. B., N. Boley, R. Eisman, G. E. May, M. H. Stoiber, *et al.*, 2014 Diversity and dynamics of the Drosophila transcriptome. Nature **512**: 393–399.

Chakraborty, M., N. W. VanKuren, R. Zhao, X. Zhang, S. Kalsow, *et al.*, 2018 Hidden genetic variation shapes the structure of functional elements in Drosophila. Nat Genet **50**: 20–25.

Lee, H., C. J. McManus, D.-Y. Cho, M. Eaton, F. Renda, *et al.*, 2014 DNA copy number evolution in Drosophila cell lines. Genome Biol **15**: R70.

Li, H., 2015 wgsim.

Li, H., B. Handsaker, A. Wysoker, T. Fennell, J. Ruan, *et al.*, 2009 The Sequence Alignment/Map format and SAMtools. Bioinformatics **25**: 2078–2079.

Linheiro, R. S. and C. M. Bergman, 2012 Whole genome resequencing reveals natural target site preferences of transposable elements in Drosophila melanogaster. PLOS One **7**: e30008.

Nelson, M. G., R. S. Linheiro, and C. M. Bergman, 2017 McClintock: an integrated pipeline for detecting transposable element insertions in whole-genome shotgun sequencing data. G3 **7**: 2749–2762.

Rahman, R., G.-w. Chirn, A. Kanodia, Y. A. Sytnikova, B. Brembs, *et al.*, 2015 Unique transposon landscapes are pervasive across Drosophila melanogaster genomes. Nucleic Acids Res **43**: 10655–10672.

Stoiber, M., S. Celniker, L. Cherbas, B. Brown, and P. Cherbas, 2016 Diverse Hormone Response Networks in 41 Independent Drosophila Cell Lines. G3 **6**: 683–694.

Yang, N., S. P. Srivastav, R. Rahman, Q. Ma, G. Dayama, *et al.*, 2021 Transposable element landscape changes are buffered by RNA silencing in aging Drosophila. bioRxiv .
